# VISUAL network analysis reveals the role of BEH3 as a stabilizer in the secondary vascular development in Arabidopsis

**DOI:** 10.1101/2021.01.19.427273

**Authors:** Tomoyuki Furuya, Masato Saito, Haruka Uchimura, Akiko Satake, Shohei Nosaki, Takuya Miyakawa, Shunji Shimadzu, Wataru Yamori, Masaru Tanokura, Hiroo Fukuda, Yuki Kondo

## Abstract

During secondary growth in plants, vascular stem cells located in the cambium continuously undergo self-renewal and differentiation throughout the lifetime. Recent cell-sorting technique enables to uncover transcriptional regulatory framework for cambial cells. However, the mechanisms underlying the robust control of vascular stem cells have not been understood yet. By coexpression network analysis using multiple transcriptome datasets of an ectopic vascular cell transdifferentiation system using Arabidopsis cotyledons, VISUAL, we newly identified a cambium-specific gene module from an alternative approach. The cambium gene list included a transcription factor BES1/BZR1 homolog 3 (BEH3), whose homolog BES1 is known to control vascular stem cell maintenance negatively. Interestingly, the vascular size of the *beh3* mutants showed a large variation, implying the role of BEH3 as a stabilizer. BEH3 almost lost the transcriptional repressor activity and functioned antagonistically with other BES/BZR members via competitive binding to the same motif BRRE. Indeed, mathematical modeling suggests that the competitive relationship among BES/BZRs leads to the robust regulation of vascular stem cells.

## Introduction

Continuous stem cell activity relies on a strict balance between cell proliferation and cell differentiation. Plant vascular stem cells located in the cambium proliferate by themselves, then continuously giving rise to conductive tissues consisting of xylem and phloem cells, as bifacial stem cells (Shi et al., 2017; De Rybel et al., 2016; Fischer et al., 2019). Previous genetic studies have indicated that vascular stem cells are maintained by cell-cell communication via a peptide hormone tracheary element differentiation inhibitory factor (TDIF) and its specific receptor PHLOEM INTERCALATED WITH XYLEM (PXY) / TDIF RECEPTOR (TDR) (Ito et al., 2006; Fisher et al., 2007; Hirakawa et al., 2008). For the maintenance of vascular stem cells, the TDIF-PXY/TDR signaling promotes cell proliferation via WUS-RELATED HOMEOBOX 4 (WOX4) and WOX14 (Hirakawa et al., 2010, Etchells et al., 2010) and represses cell differentiation via BRI1-EMS-SUPPRESSOR 1 (BES1) (Kondo et al., 2014). In addition, the TDIF-PXY/TDR signaling genetically cross-talks with several peptide or hormonal signaling pathways, thereby forming a complicated regulatory network (Han et al., 2018; Etchells et al., 2012; Kondo and Fukuda, 2015). As another attempt to identify stem cell regulators, transcriptome analysis using fluorescence activated cell sorting (FACS) and fluorescence activated nuclear sorting (FANS) method has been conducted. FACS transcriptome analysis with a (pro)cambium marker *ARABIDOPSIS RESPONSE REGULATOR 15* (*ARR15*) *promoter:GFP* educed a list of cambium-expressed transcription factors, including *AINTEGUMENTA* (*ANT*), *KNOTTED-like from Arabidopsis thaliana 1* (*KNAT1*), *LOB DOMAIN-CONTAINING PROTEIN 3* and *4* (*LBD3* and *LBD4*), *SHORT VEGETATIVE PHASE* (*SVP*), and *PETAL LOSS* (*PTL*) (Zang et al. 2019). Furthermore, in the inflorescence stem, FANS transcriptome analysis using tissue-specific promoters including *PXY/TDR* promoter for proximal cambial cells and *SUPPRESSOR OF MAX2 1-LIKE PROTEIN 5* (*SMXL5)* promoter for distal cambial cells provided informative vascular tissue-specific gene expression profiles (Shi et al., 2020). Nevertheless, the molecular mechanism enabling a robust control of vascular stem cells remains largely unknown.

The vascular system consists of various types of vascular cells at different developmental stages, preventing the sequential analysis of vascular development with a focus on bifacial stem cells. To simply understand a complex biological phenomenon such as vascular development, reconstitutive approach has great experimental advantages with regard to molecular genetic studies (Kondo 2018). We previously established a tissue culture system, Vascular cell Induction culture System Using Arabidopsis Leaves (VISUAL), which can synchronously and efficiently induce xylem and phloem cells from mesophyll cells via vascular stem cell (Kondo et al., 2016). The loss-of-function mutant for the phloem regulator ALTERED PHLOEM DEVELOPMENT (APL) lacks phloem sieve element differentiation. In addition to the microarray data with the *apl* mutants, the datasets of the FACS with a phloem marker gene, *SIEVE ELEMENT OCCLUSION RELATED 1* (*SEOR1*) in VISUAL enabled the construction of a coexpression gene network from early to late phloem sieve element differentiation process (Kondo et al. 2016). Although such a network analysis is useful for temporal dissection of vascular differentiation process, this gene network is still only limited to phloem sieve element differentiation. Recently, we discovered that the loss-of-function *bes1* mutants showed a dramatic impairment in the differentiation of xylem and phloem cells, resulting in the accumulation of vascular stem cells in the VISUAL (Kondo et al., 2014; Saito et al., 2018; Kondo et al., 2015). Then we further integrated *bes1* transcriptome and time-course (every 6 hours) transcriptome datasets in order to visualize the gene network for the entire vascular differentiation process.

In this study, we newly established a coexpression network of the VISUAL differentiation process utilizing multiple transcriptome datasets including the *bes1* mutants. As a result, we succeeded in the identification of gene modules corresponding to not only phloem but also xylem and cambial cells. These classifications were validated with a comparison to the already-existing vascular FACS and FANS datasets. Through the network analysis, one of BES/BZR homolog BEH3 was found as a potential vascular stem cell regulator. Indeed, a knock-out mutant of BEH3, *beh3-1*, exhibited a high variation in vascular size among individual samples, suggesting that BEH3 functions as a stabilizer of vascular stem cells. Further genetic and molecular studies revealed that BEH3 has an opposite function toward other BES/BZR homologs by interfering with other BES/BZR homologs via competitive binding to the BR response element (BRRE). Moreover, mathematical modeling analysis showed that the competitive relationship among BEH3 and other BES/BZR homologs enables the robust control of vascular stem cells.

## RESULTS

### Coexpression network analysis using the multiple VISUAL transcriptome datasets identifies a cambium-related gene module

To find out novel regulators for vascular stem cells, we conducted a gene network analysis using the transdifferentiation system VISUAL (Figure 1A; Kondo et al., 2015). First, we selected 855 VISUAL-induced genes that were up-regulated highly (more than eight-fold) in microarray data (0 h vs 72 h after VISUAL induction) (Figure 1B; Supplemental Data set 1). Since the selected 855 VISUAL-induced genes should include xylem-, phloem- and cambial-related genes, we constructed a coexpression gene network for these genes with the use of multiple VISUAL transcriptome datasets including (1) 6-h interval time-course, (2) cell-sorting with the phloem marker *pSEOR1:SEOR1-YFP*, (3) phloem deficient *apl* mutants, and (4) *bes1-1* mutants (Figure 1A). Here the *bes1-1* mutant transcriptome datasets were used for dissecting cambial cells and differentiated xylem/phloem cells, because the *bes1-1* mutants significantly accumulate cambial cells due to the inhibition of xylem and phloem differentiation in the VISUAL (Figure 1A). Coexpression network using the WGCNA package divided the 855 VISUAL-induced genes into four distinct modules (Module-I to IV) and 11 outgroup genes (Others, O) (Figure 1C). Expression of Module-III and -IV genes were down-regulated in the *bes1-1* mutants (Figure 1C), suggesting that these modules were considered as differentiated xylem and phloem cells-related ones. In addition, expression of Module-III genes was high in the SEOR1-positive cell-sorting datasets and was down-regulated in the *apl* mutants, whereas expression of Module-IV genes was unchanged in the cell-sorting and the *apl* mutant datasets (Figure 1D), suggesting that Module-III and -IV represents phloem-related and xylem-related genes, respectively. Consistent with this idea, the known phloem-related genes such as *RAB GTPASE HOMOLOG C2A* (*RABC2A*), *BREVIS RADIX* (*BRX*), *GLUCAN SYNTHASE-LIKE 7* (*GSL07*), and *NAC DOMAIN CONTAINING PROTEIN 45* (*NAC045*), were found in Module-III, whereas xylem-related genes such as *IRREGULAR XYLEM 3* (*IRX3*), *XYLEM CYSTEINE PEPTIDASE 1* (*XCP1*), and *VASCULAR-RELATED NAC-DOMAIN 6* (*VND6*), were in Module-IV (Figure 1C). On the other hand, expression of Module-I and -II genes was induced at earlier timing and was remained higher even in the *bes1-1* mutants than that of Module-III and -IV genes (Figure 1B and 1D), suggesting that Module-I and -II genes correspond to (pro)cambial-related genes. To validate these characterizations, we investigated gene expression of each module in the transcriptome datasets of FACS in roots and FANS in stems obtained using the cell-type specific markers (Brady et al. 2007, Zang et al. 2019, Shi et al. 2020). Expectedly, expression of Module-III and -IV genes were quite high in the phloem specific *APLpro* datasets and the xylem specific *S18* and *VND7pro* datasets, respectively (Figure 1E; Supplemental Figure 1A). As for Module-I and II, the genes in Module-I were relatively highly expressed in the procambium (ARRIII ± BAP) and expression of Module-II gene is more specific to the developing cambium and cambium (ARRI ± BAP and ARRII ± BAP) in the FACS datasets (Figure 1E). These results suggest that Module-I and Module-II represents procambium-related and cambium-related genes, respectively. Consistent with this idea, expression of Module-I genes was expressed much earlier than that of Module-II genes (Figure 1B).

**Figure 1.**
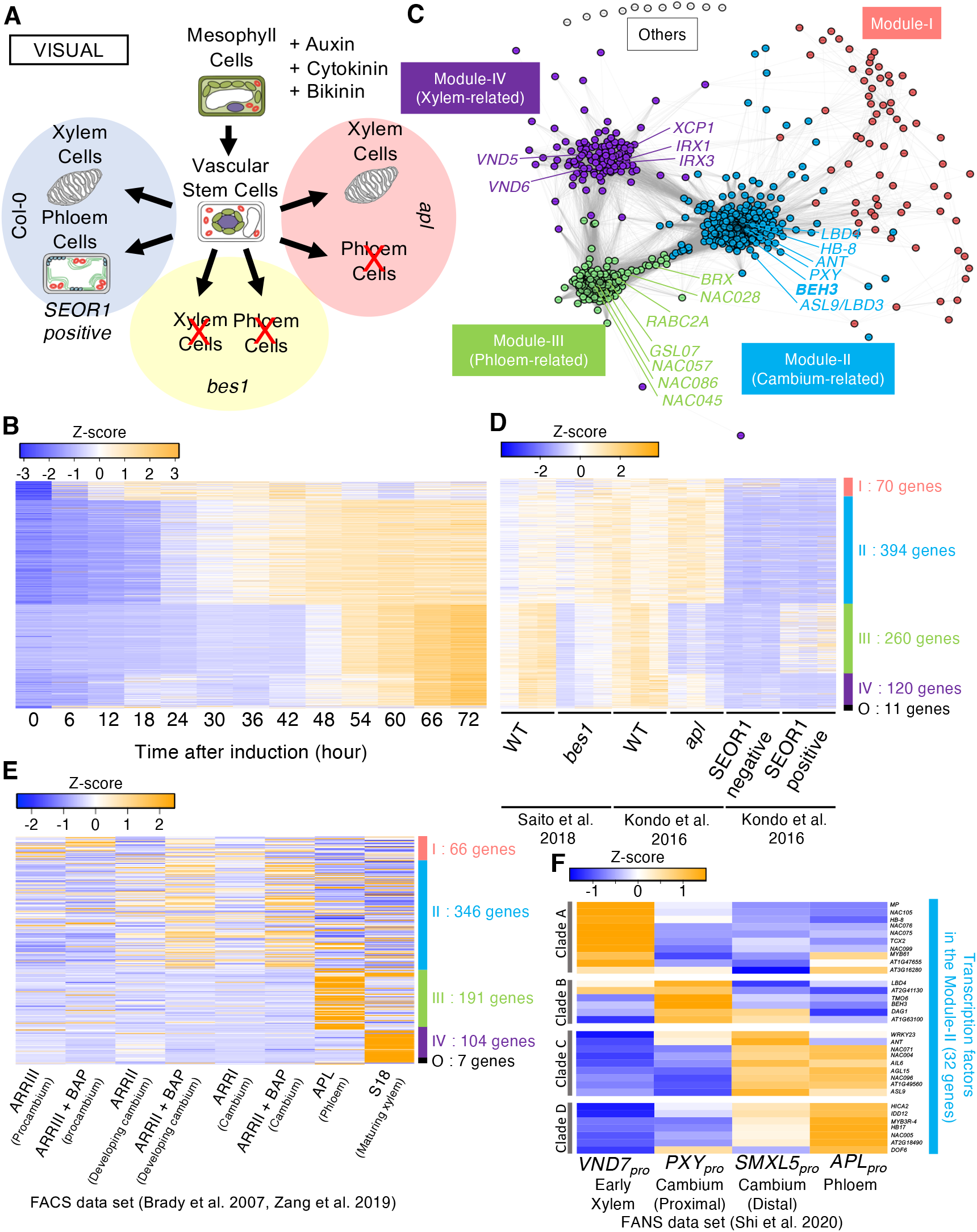
Classification of the VISUAL-induced genes based on the gene coexpression network analysis. **(A)** Schematic showing the transdifferentiation process in Col-0 (WT, left), *bes1* mutants (lower), and *apl* mutants (right). **(B)** Expression profiles of the VISUAL-induced genes in the VISUAL time-course microarray data. The VISUAL-induced genes were defined as >8-fold changed genes when compared VISUAL at 0 and 72 h. Expression was visualized with a heat map image according to Z score according to their expression levels. Color bars in the right side indicates each module in accordance with (C). **(C)** Coexpression network of the VISUAL-induced genes using the WGCNA package. Four distinct modules are highlighted with different colors. A node represents a gene. An edge indicates high correlation between nodes (TOM > 0.02). **(D)** Expression profiles of the VISUAL-induced genes in the multiple VISUAL microarray datasets. VISUAL microarray datasets were provided from each comparison between WT and *bes1-1* (Saito et al. 2018), WT and *apl* (Kondo et al. 2016), and SEOR1-negative and SEOR1-positive cells (Kondo et al. 2016). **(E)** Expression profiles of the VISUAL-induced genes in the FACS dataset (Bardy et al., 2007, Zang et al., 2019). **(F)** Expression profiles of the cambium-related genes (Module-II) in the FANS dataset (Shi et al., 2020). The gene names are shown on the right side.

We further analyzed the Module-II genes for analyzing stem cell regulation. The Module-II consists of a total of 394 genes including some known cambium-related genes, such as *PYX/TDR, SMXL5, ANT, LBD3, LBD4*, and *HOMEOBOX GENE 8* (*HB-8*) (Figure 1B – 1D). Of them, 32 genes encoding transcription factors were investigated in more detail (Supplemental Data set 2). Previous studies have revealed that vascular stem cells locate in the overlapped region of *PXY/TDR* and *SMXL5* expression, which marks proximal/xylem and distal/phloem side of cambium, respectively (Shi et al. 2019). According to the FANS transcriptome datasets (Shi et al., 2020), 13 out of the 32 transcription factor genes are expressed mainly in *PXY/TDRpro* and/or *SMXL5pro* (Figure 1F). They included not only known cambial regulators such as *ANT* (Randall et al., 2015) and *LBD4* (Zang et al., 2019; Smit et al., 2020), but also novel candidates such as *BEH3, DOF AFFECTING GERMINATION 1* (*DAG1*), and *At1g63100* (Figure 1E). The analysis of these newly selected genes may provide a novel insight into regulation of cambial activity.

### BEH3 stabilizes secondary vascular development

We recently showed that BES1 and its closest homolog BRASSINAZOLE RESISTANT1 (BZR1) redundantly function to promote vascular stem cell differentiation in VISUAL (Saito et al., 2018). Therefore, we chose BEH3 as a potential vascular stem cell-specific regulator for further studies. Among 6 BES/BZR homologs in *Arabidopsis thaliana*, only *BEH3* was found in the cambium-related module. Indeed, only *BEH3* expression levels were remarkably increased in the time course RNA-seq data of the VISUAL (Figure 2B), and the GUS expression domain of *pBEH3:GUS* gradually expanded over time (Figure 2C). Consistent with the observation in the VISUAL, *pBEH3:GUS* was preferentially expressed in the vascular tissues including cambium of leaves and hypocotyls (Figure 2D and 2E). Then to investigate the function of BEH3 in vascular development, we analyzed the BEH3 knock-out mutant, *beh3-1*, which has a T-DNA insertion on its 1st exon (Lachowiec et al., 2018). Here we compared vascular secondary growth in the WT and the *beh3-1* mutants grown under short-day condition. The area inside the cambium in the *beh3-1* mutants were slightly but significantly decreased when compared with the WT (Figure 2F and 2G; Supplemental Figure 2). In addition, we realized that the *beh3-1* mutants seem to have a higher variation in area inside the cambium than the WT (Figure 2H). Indeed, the values of coefficient of variation (CV) between the *beh3-1* mutants and WT plants were significantly different in two statistical analyses: asymptotic and Modified signed-likelihood ratio test (M-SLRT) (Feltz and Miller, 1996; Krishnamoorthy and Lee, 2014) (Figure 2I). These results suggest that the presence of BEH3 contributes to not only promoting but also stabilizing secondary vascular development.

**Figure 2.**
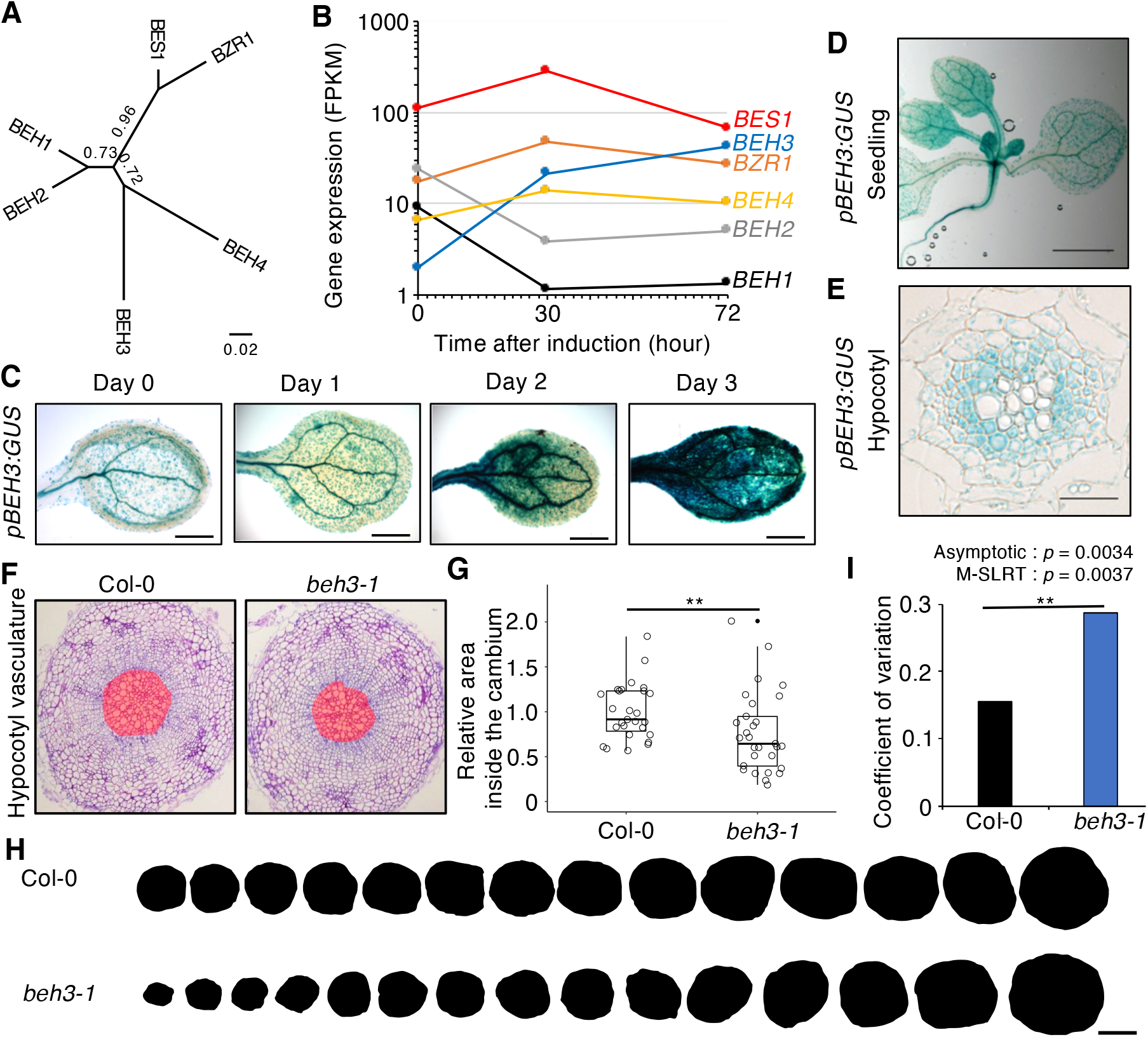
BEH3 provides the stability for secondary growth in the cambium. **(A)** An unrooted phylogenetic tree of Arabidopsis BES/BZR family members. **(B)** Time-course expression analysis of BES1 homologs genes using VISUAL. Gene expression levels at each time point were analyzed by RNA-seq. **(C-E)** GUS activity in *pBEH3:GUS* transgenic plants. GUS staining was performed using cotyledons subjected to VISUAL (C) shoot of 20-day-old seedling (D) and hypocotyl section of 7-day-old seedling (E). **(F)** Sections of hypocotyl vasculature of WT and *beh3-1* mutant plants grown under short-day conditions for 21 days. **(G)** Relative areas inside the cambium were calculated in comparison with the mean value of the WT (\*\**P* < 0.01; Student’s *t*-test, *n* = 28 [WT], 30 [*beh3-1*]). **(H)** Size variation inside the cambium of WT and *beh3* plants. Black circles indicate areas inside the cambium of each individual for the odd number samples in ascending order of (F). **(I)** CV of the areas inside the cambium of WT and *beh3* plants. Statistical analysis of differences between the WT and *beh3* mutants using Asymptotic and M-SLART. Scale bars = 1 mm (C), 2 mm (D), and 100 µm (E,F,I).

### BEH3 functions oppositely to BES1 in vascular development

In *Arabidopsis thaliana*, 6 BES/BZR transcription factors have the redundant function in brassinosteroid (BR) signaling (Yin et al., 2005; Lachowiec et al., 2018; Chen et al., 2019a; Chen et al., 2019b; Nolan et al., 2020). As previously reported (Kondo et al., 2014; Saito et al., 2018), *bes1-D*, which has a mutation in protein degradation PEST sequence, exhibited a reduced number of (pro)cambial cell layers between differentiated xylem (red) and phloem (green) in hypocotyls (Figure 3A). To compare the function of BEH3 and BES1, vascular phenotype of gain-of-function mutant for BEH3 was analyzed. Since BEH3 does not possess any obvious PEST domain, we generated a transgenic stable line of β-estradiol-inducible BEH3 overexpressor (*pER8:BEH3-CFP*). In contrast to the *BES1* gain-of-function mutants, *BEH3* overexpression lines showed an increase in the number of (pro)cambium layers (Figure 3B). In association with this tendency, the vascular area was also enlarged in the two independent *BEH3* overexpression lines (Figure 3C and 3D; Supplemental Figure 3), which is not contradictory to the *beh3-1* mutant phenotype. These results suggest that BEH3 has an opposite function to BES1 during secondary vascular development.

**Figure 3.**
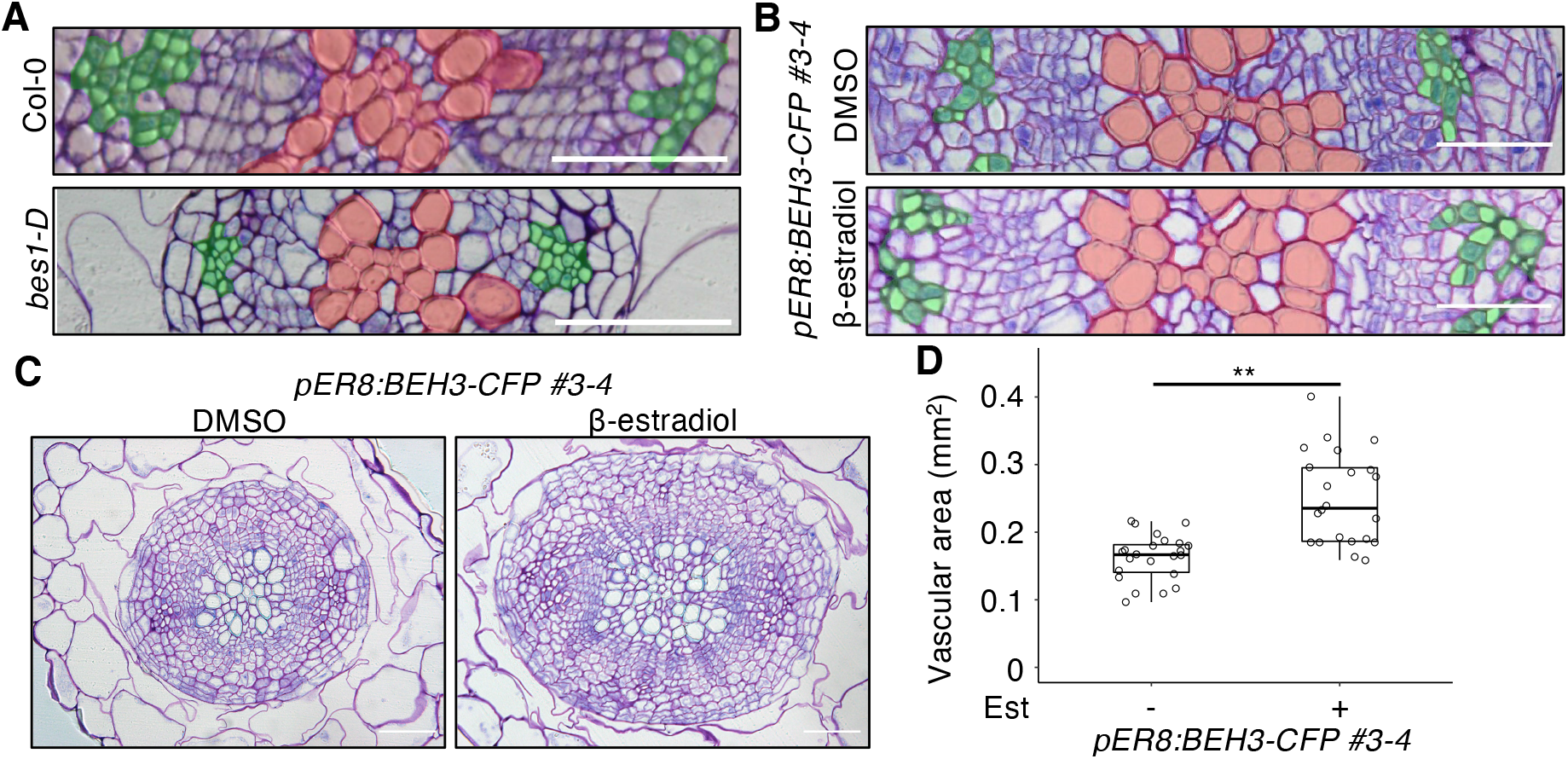
BEH3 overexpression promotes secondary growth. **(A)** Sections of hypocotyl vasculature of 11-day-old WT and *bes1-D* seedlings. Red and green area indicates xylem cells and phloem cells, respectively. **(B-C)** Sections of hypocotyl vasculature of β-estradiol-inducible BEH3-overexpressing plants treated with (+) or without (-) 5 µM β-estradiol for 12 days. Red and green area in (B) indicate xylem cells and phloem cells, respectively. **(D)** Calculated vascular areas in (C) are shown using box-and-whisker plots. White circles indicate values of each sample (\**P* < 0.05, \*\**P* < 0.01; Student’s *t*-test; *n* = 23). Scale bars = 100 µm (A–C).

As described above, the *bes1-1* mutants disrupt cell differentiation from vascular stem cells in the VISUAL (Figure 1A). To investigate the function of BEH3 and other BES/BZR homologs, we evaluated the contribution of each BES/BZR homolog to xylem differentiation with a semi-automatic calculation method using a xylem indicator, BF-170 (Nurani et al., 2020) in VISUAL (Figure 4A and 4B; Supplemental Figure 4). As for the analysis with single mutants, *beh4-1* as well as *bes1-1* and *bzr1-1* showed reduced xylem differentiation ratio in the VISUAL compared with the WT (Fig. 4C and 4D; Supplemental Figure 5), indicating that BEH4 also promotes xylem differentiation. On the other hand, xylem differentiation ratio in *beh3* as well as *beh1* and *beh2* were comparative to the WT (Figure 4B and 4C). Next, we examined the impact of mutating a BES1 homolog in the *bes1-1* mutant background as double mutants. Only the *bes1-1 beh3-1* double mutants showed a high xylem differentiation ratio, similar to that observed in the WT, whereas none of the other mutants rescued the *bes1* phenotype (Figure 4B and 4C; Supplemental Figure 5 and 6). The *bes1* mutants disrupt the differentiation from vascular stem cells into not only xylem but also phloem cells (Saito et al., 2018), accompanied by the decrease in expression levels of xylem-specific (*IRX3*) and phloem-specific (*SEOR1*) marker genes (Figure 4E). This reduction in the *bes1-1* mutants was recovered up to WT levels by the additional *beh3-1* mutation (Figure 4E). Quantitative expression analysis also indicated that the single *beh3-1* mutants slightly increase the expression levels of *IRX3* and *SEOR1* at 48 h, when compared to the WT (Figure 4E; Supplemental Table 1). These genetic data suggest that BEH3 functions oppositely to BES1 also in the VISUAL.

**Figure 4.**
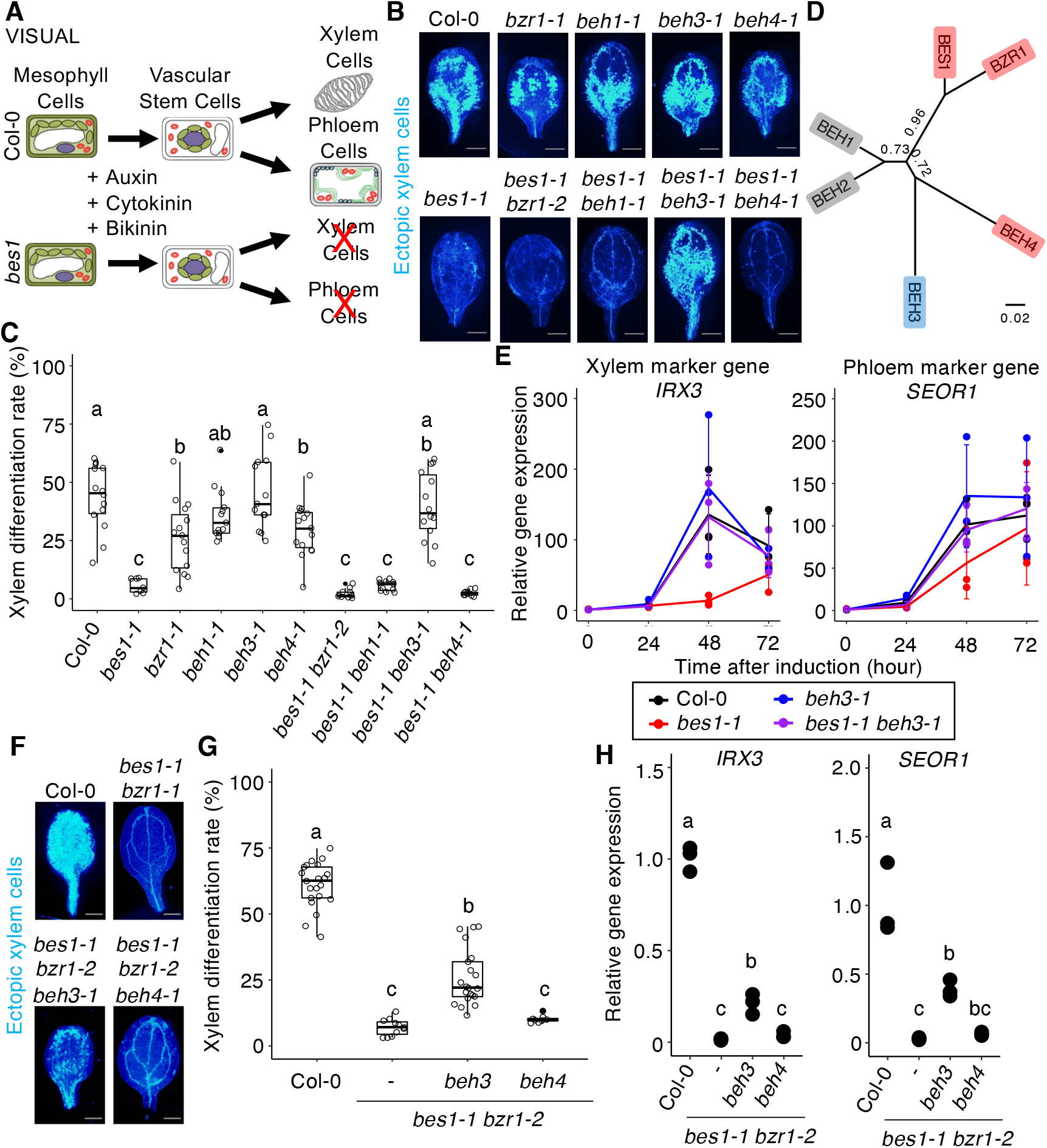
BEH3 antagonistically functions against BES1 and BZR1 in VISUAL. **(A)** Schematic showing the transdifferentiation process in Col-0 (WT, upper) and *bes1* mutants (lower). **(B, C)** Ectopic xylem cell differentiation ratio in a combination of *bes/bzr* mutants. Bright blue signal in (B) indicates xylem cell wall stained with BF-170. Quantified xylem differentiation ratio is shown using box-and-whisker plots in (C). White circles indicate the xylem differentiation ratio for each sample. Different letters indicate significant differences (*P* < 0.05; Tukey-Kramer test). **(D)** Effects of BES/BZR family members toward vascular cell differentiation in VISUAL. Red, blue, and grey indicate “promote”, “suppress”, and “no effect”, respectively. **(E)** Time-course expression analysis of xylem-specific (*IRX3*) and phloem-specific (*SEOR1*) marker genes. Relative fold changes were calculated in comparison with the WT at 0 h after induction. Data represent mean ± standard deviation (SD; n = 3). Results of two-way analysis of variance (ANOVA) are summarized in Supplemental Table 1. **(F, G)** Ectopic xylem differentiation in triple mutants. Bright blue signal in (F) indicates xylem cell wall stained with BF-170. Xylem differentiation ratio is shown using box-and-whisker plots in (G). **(H)** Expression levels of *IRX3* and *SEOR1* in triple mutants at 48 h after induction. Relative fold changes were calculated in comparison with the WT at 0 h after induction (*n* = 3). Different letters indicate significant differences (*P* < 0.05; Tukey-Kramer test). Scale bars = 1 cm in (B,F).

**Figure 5.**
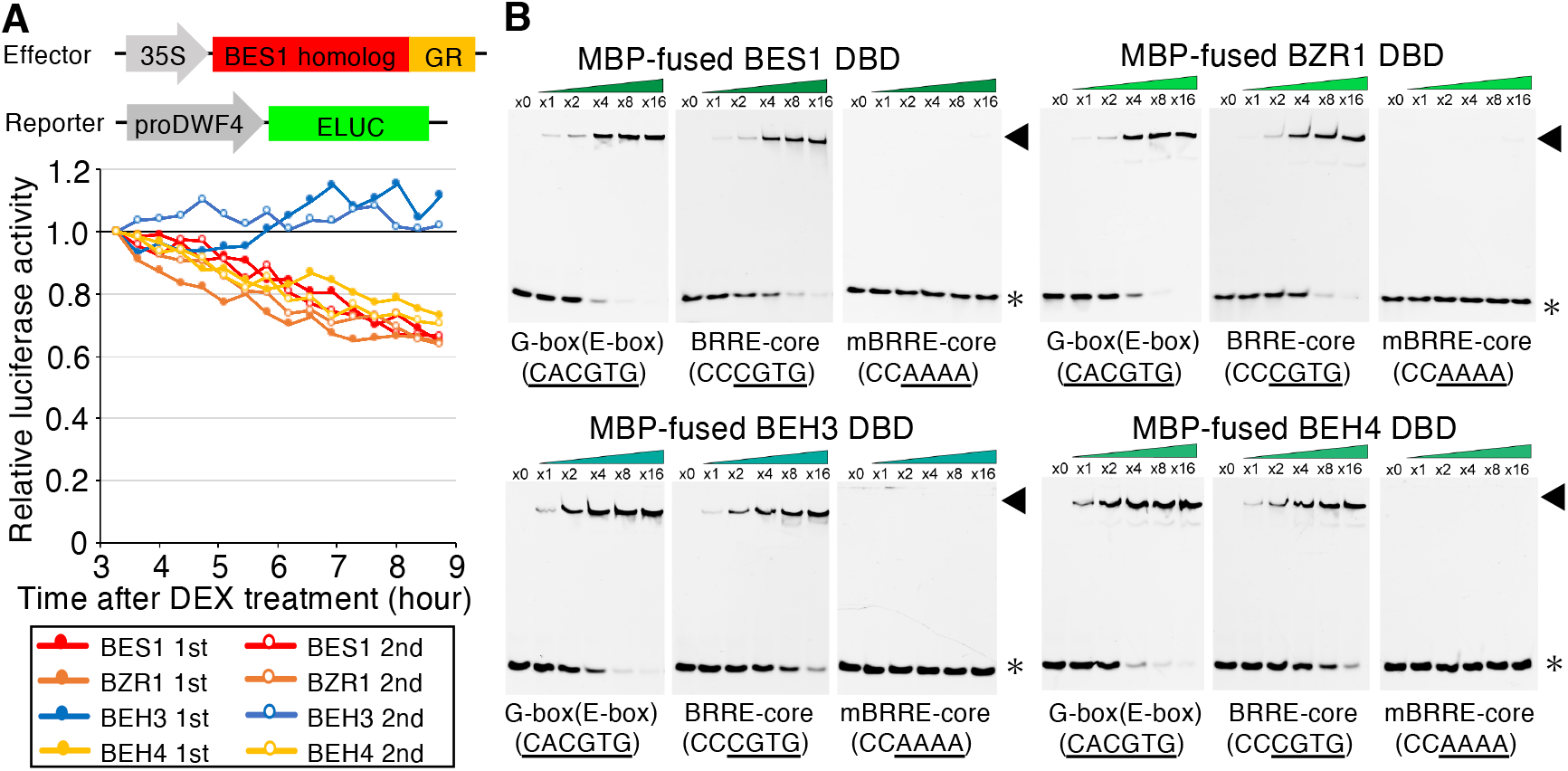
BEH3 competes with other BES/BZR homologs. **(A)** Transcriptional repressor activity of BES/BZR proteins in the *Nicotiana benthamiana* transient expression assay. Luciferase (LUC) activities in *pDWF4:ELUC* transgenic plants were calculated as average photon counts per second of six leaf disks and were normalized relative to values of DEX-treated samples harboring an empty effector construct. **(B)** DNA-binding ability of BES/BZR proteins in the electrophoretic mobility shift assay (EMSA). Asterisks and arrowheads indicate the positions of free DNA and DNA-binding domain (DBD)-containing protein-bound DNA, respectively.

**Figure 6.**
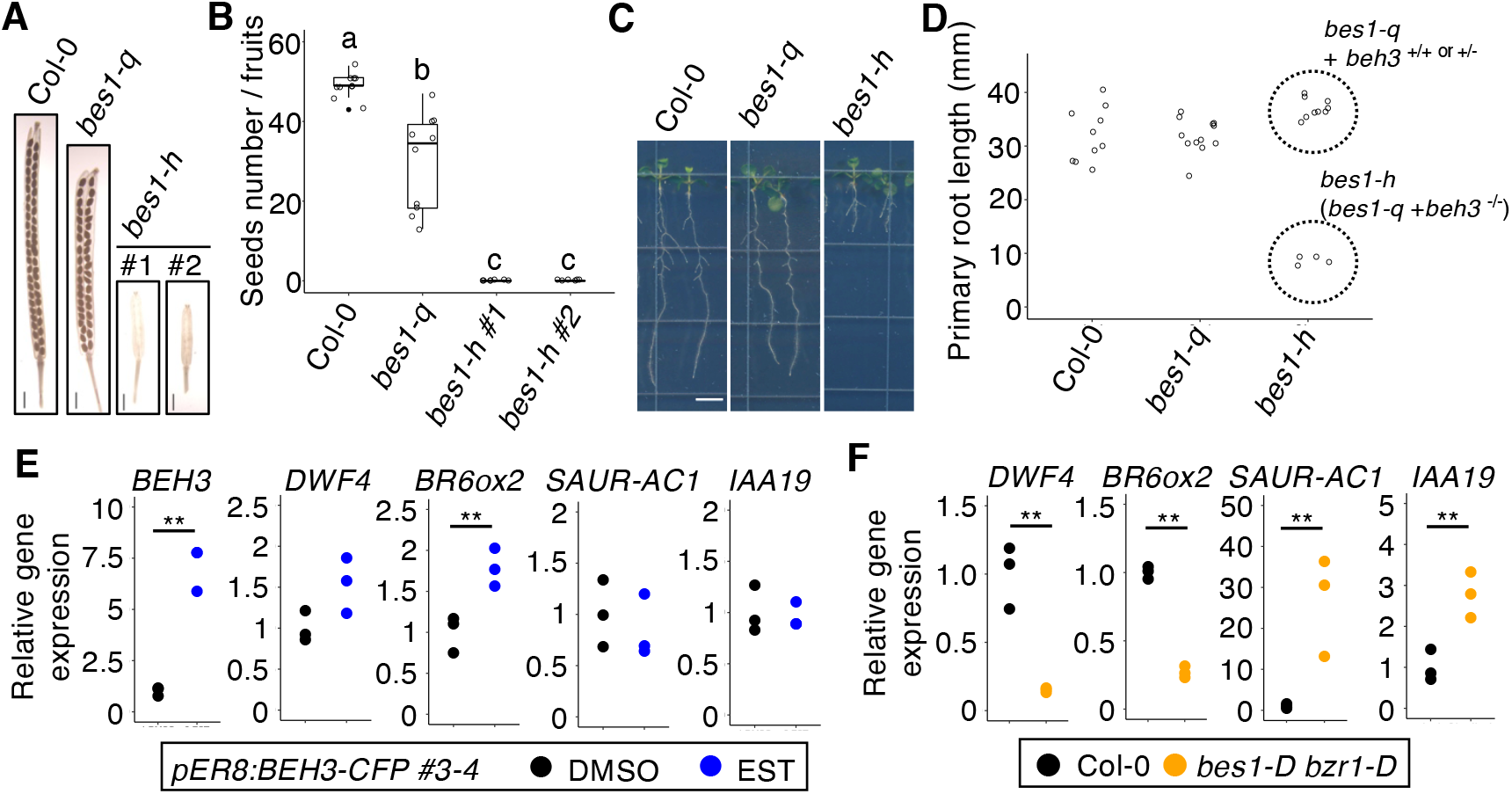
BEH3 passively inhibits the repressor activity of other BES/BZR homologs. **(A, B)** Development of fruits and seeds in *bes1* quintuple (*bes1-q*) and hextuple (*bes1-h*) mutants. Number of seeds in each mature fruit are shown using box-and-whisker plots in (B). Different letters indicate significant differences (*P* < 0.05; Tukey-Kramer test; *n* = 10). **(C, D)** Primary root length in 11-day-old *bes1-q* and *bes1-h* mutant seedlings. To obtain homozygous *bes1-h* mutant plants, self-fertilized progenies of the T2 heterozygous *bes1-q* mutants, which containing *beh3*^*+/-*^ mutation, were used. *n* = 10 (WT), 12 (*bes1-q*), and 13 (*bes1-h*). **(E)** Expression levels of BES/BZR target genes in the estradiol-inducible BEH3 overexpression line. Eight-day-old seedlings were treated with 10 µM β-estradiol for 9h. Relative gene expression levels were calculated in comparison with the DMSO-treated sample (\*\**P* < 0.01; Student’s *t*-test; *n* = 3). **(F)** Expression level of BES/BZR target genes in *bes1-D bzr1-D*. Nine-day-old seedlings were used. Relative gene expression levels were calculated in comparison with the WT (\*\**P* < 0.01; Student’s *t*-test; *n* = 3). Scale bars = 1 mm (A and C).

### BEH3 has a competitive relationship with other BES/BZR homologs

To unravel the molecular basis of the antagonistic function of BEH3, we examined the functional difference among BEH3 and other BES/BZR proteins. Amino acid sequence alignments suggested no obvious difference in the known functional domains/motifs, such as DNA-binding domain (DBD) and EAR motif (Yin et al., 2002; Wang et al., 2002; Oh et al., 2014). It is known that BES1 and BZR1 repress the transcription of a BR biosynthetic gene, *DWARF4* (*DWF4*), by directly binding to the BRRE and/or E-box motif in the *DWF4* promoter (He et al., 2005; Sun et al., 2010; Yu et al., 2011; Chung et al., 2011; Nosaki et al., 2018). Therefore, we tested the transcriptional control activity of BES/BZR members against the *pDWF4: Emerald Luciferase (ELUC)* construct transiently expressed in *Nicotiana benthamiana* leaves. The signal intensity of ELUC gradually decreased from 3 to 9 h after activating BES1, BZR1, and BEH4 by induction system using dexamethasone (DEX) and rat glucocorticoid receptor (GR) (Aoyama et al., 1995), whereas BEH3 activation did not alter ELUC activity (Figure 5A). Electrophoretic mobility shift assay (EMSA) for the BRRE core motif and E-box motif with DBD of BES/BZR members indicated that BEH3 can bind to the BRRE and E-box, similar to BES1, BZR1, and BEH4 (Nosaki et al., 2018) (Figure 5B; Supplemental Figure 7). These results suggest that BEH3 binds to the BRRE and E-box motif but exhibits much lower transcriptional repressor activity than other BES/BZR homologs. When BEH3 and BES1 were simultaneously expressed in *N. benthamiana* leaves, repressor activity of BES1 against *pDWF4:ELUC* was weakened (Supplemental Figure 8). By contrast, co-expression of BZR1 with BES1 did not show such a combinatory repressive effect (Supplemental Figure 8), suggesting the possibility that BEH3 interferes with other BES/BZR homologs via the binding competition.

**Figure 7.**
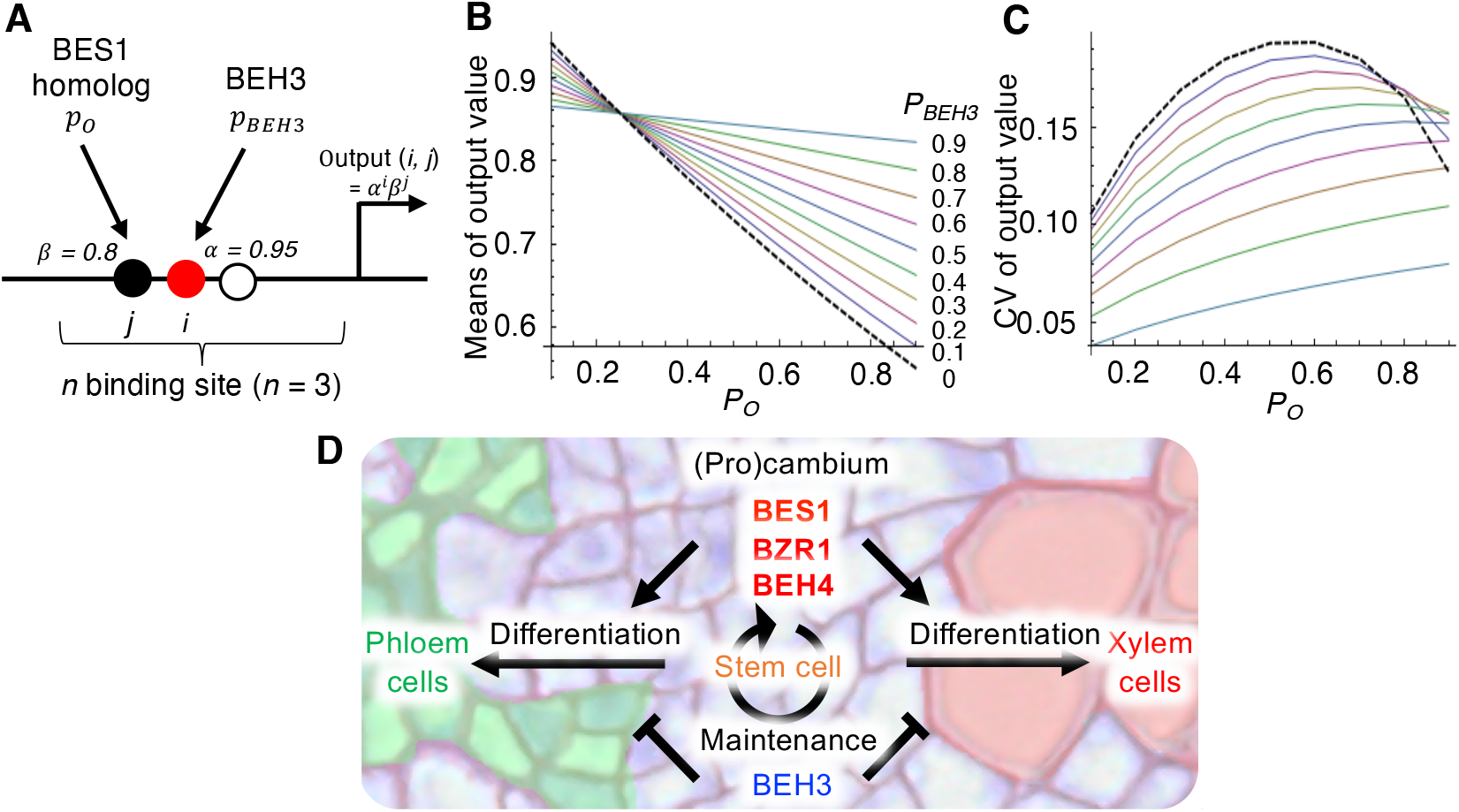
Competitive relationship among BES/BZR family provides the robustness for the control of vascular stem cell activity. **(A)** Schematic representation of the mathematical model that describes competitive binding between BEH3 and other homologs. BEH3 binds to i of n binding sites at a certain binding rate (*P*_*BEH3*_). Then, other BES1 homologs (*P*_*o*_) bind to *j* of *n-i* binding sites at a certain binding rate. Signaling output (*i, j*) is defined as *α*^*i*^β^*j*^; αand β represent coefficients of transcriptional activity (<1 in the case of a repressor) of BEH3 and other BES1 homologs, respectively. **(B, C)** Simulated signaling output when verifying both *P*_*BEH3*_ and *P*_*o*_ (*n* =3; α= 0.95; β = 0.8). Means and c oefficient of variation (CV) (CV = standard deviation/mean) of output values are shown in (B) and (C), respectively. **(D)** Model showing the role of BES/BZR family members in the robust control of vascular stem cell differentiation and maintenance. Scale bars = 100 µm (H–J).

In the VISUAL, the mutation in *BES1* did not affect *BEH3* expression, and the mutation in *BEH3* did not affect *BES1* expression in VISUAL (Supplemental Figure 9), indicating that there seems not to be an epistatic relationship between *BEH3* and *BES1*. Then to genetically investigate the competitive relationship among *BEH3* and other *BES/BZR* homologs, several combinations of triple mutants were analyzed in VISUAL. Although the *beh3-1* mutant allele could completely rescue the phenotype of the *bes1-1* single mutants (Figure 4B – 4D), restoration of xylem differentiation by the *beh3-1* mutant allele was only partial in the background of *bes1-1 bzr1-2* double mutants (Figure 4F and 4G). Such a partial rescue was not detected in the *bes1-1 bzr1-2 beh4-1* triple mutants at all (Figure 4F and G). Consistent with the xylem differentiation ratio, the *bes1-1 bzr1-2 beh3-1* triple mutants were intermediate between WT and the *bes1-1 bzr1-2* double mutants in terms of expression levels of xylem and phloem marker genes (Figure 4H). Taken together, these results suggest that BEH3 acts antagonistically with other homologs in a competitive manner in the VISUAL.

### Competitive inhibition by BEH3 may be limited to the transcriptional repressor activity of other BES/BZR homologs

Previous studies showed that a *bes1* hextuple mutants, in which all BES/BZR members are knocked-out, exhibits male sterility (Chen et al., 2019a). If BEH3 always has a competitive role against other BES/BZR homologs, *bes1 bzr1 beh1 beh2 beh4* quintuple mutants are expected to show the same phenotype as *bes1 bzr1 beh1 beh2 beh3 beh4* hextuple mutants. Then we generated a *bes1* hextuple mutants (*bes1-1 bzr1-2 beh1-2 beh2-5 beh3-3 beh4-1*) and a *bes1* quintuple mutants (*bes1-1 bzr1-2 beh1-2 beh2-5 beh4-1*) using CRISPR/Cas9 system. Consistent with the previous reports, the newly established *bes1* hextuple mutants showed shorter siliques with no seed (Figure 6A and 6B; Supplemental Figure 10). However, the *bes1* quintuple mutants successfully produced seeds in siliques (Figure 6A and 6B). In addition, while the primary root length of the *bes1* hextuple mutants were shorter than the WT, that of the *bes1* quintuple mutants were comparable to the WT (Figure 6C and 6D). These results suggest that BEH3 partially remains the same function to other BES/BZR homologs.

BES1 and BZR1 is known to function as both transcriptional activators and repressors (Yin et al., 2005; He et al., 2005; Nolan et al., 2020). Then, we investigated to which extent BEH3 affects the expression of known BES/BZR-target genes in planta. Here we used the β-estradiol-inducible *BEH3* overexpressor *pER8:BEH3-CFP*, in which expression of *BEH3* was increased upon β-estradiol treatment (Figure 6E; Supplemental Figure 11). *DWF4* and *BRASSINOSTEROID-6-OXIDASE 2* (*BR6ox2*), as BES1-direct repressed genes (He et al., 2005; Sun et al., 2010; Yu et al., 2011), were remarkably repressed in the constitutively active mutant for *BES1* and *BZR1* (*bes1-D bzr1-D*) (Figure 6F), whereas *pER8:BEH3-CFP* transgenic plants significantly increased the mRNA levels of both *DWF4* and *BR6ox2* upon β-estradiol treatment (Figure 6E; Supplemental Figure 11; Supplemental Table 2). These results suggest that the non-canonical BEH3 can passively inhibit the transcriptional repressor activity of endogenous BES1 homologs by competing for the same binding motif. However, *BEH3* overexpression did not suppress mRNA levels of BES1-direct activated genes, *SAUR-AC1* and *INDOLE-3-ACETIC ACID INDUCIBLE 19* (*IAA19*) (Yin et al., 2005; Sun et al., 2010; Oh et al., 2012) (Figure 6E), whose expression was highly induced in the *bes1-D bzr1-D* double mutants (Figure 6F). Together with the genetic results of the *bes1* high-order mutants, our results suggest that competitive relationship among BES/BZR members may be limited mainly to their transcriptional repressor activity.

### BEH3 contributes to increasing robustness via the competition with other homologs

Our observation revealed that the *beh3* mutants increased the variation in the size of vascular area (Figure 2). Then to understand the significance of the competitive relationship between BEH3 and other BES/BZR members in phenotypic variation, a mathematical model for competitive binding was constructed with a focus on the transcriptional repressor activity. In this mathematical modeling, we set the repressor activity of BEH3 and other homologs at 0.95× and 0.8×, respectively, when they occupied one binding site in the target gene promoter. Here, we assumed three binding sites in the putative promoter and predicted the output as vascular cell number, when the binding rate of BEH3 (*P*_*BEH3*_) and other homologs (*P*_*O*_) was varied (Figure 7A). When *P*_*BEH3*_ was increased, the average predicted outputs went up because the accessibility of the binding sites for other homologs was decreased (Figure 7B). This competitive modeling can account for the phenotype that *BEH3* overexpression leads to the enlargement of vascular area (Figure 3C and 3D). On the other hand, further modeling revealed that a reduction in *P*_*BEH3*_ decreased the means of output value, which corresponds to smaller area inside the cambium in the *beh3-1* mutants when compared with the WT (Figure 2F - 2H). Moreover, because the *beh3-1* mutants exhibited higher variation in the vascular size than the WT (Figure 2F - 2H), we also simulated the shift of the CV when the value of *P*_*BEH3*_ was decreased. As a result, a reduction in *P*_*BEH3*_ increased the CV of the output (Figure 7C), which mimics the vascular phenotype of the *beh3-1* mutants. Collectively, our results indicate that the cambium-expressed BEH3 competitively inhibits transcriptional repressor activity of other BES/BZR homologs, thereby minimizing the fluctuation of the vascular stem cell activity.

## DISCUSSION

In this study, we constructed the gene coexpression network during the VISUAL differentiation process (Figure 1). With the help of the existing FACS and FANS transcriptome datasets, our network analysis categorized vascular-related genes into four distinct modules; procambium (Module-I), cambium (Module-II), phloem (Module-III) and xylem (Module-IV). Thus, reconstitute approach using the VISUAL is effective to dissect a hierarchical process of vascular cell differentiation and helpful for isolating genes involved in this process. Although the FACS datasets (Zang et al. 2019) almost share a similar tendency with our network prediction (Figure 1; Supplemental Figure 1), the overlapping rate of cambial genes was not so high (12.9 %, 45 genes/394 genes of Module-II). The low rate may result from differential cell state of cambium/vascular stem cells between “*in vivo*” and “*in vitro*” and/or technical differences between cell-sorting and whole tissue sampling. Therefore, combinatory approach by bioinformatics will allow the accurate and efficient identification of gene-of-interest expressed in the vasculature, which is usually masked by its deep location. Indeed, we successfully identified a novel cambium regulator BEH3 in the gene lists combined the Module-II genes with the FANS datasets (Shi et al., 2020). Also in the xylem and phloem development, there remain so many unidentified genes. Moreover, it is poorly understood how the cambium is established after procambium development. Therefore, our coexpression network will be informative for further studies on the sequential vascular development.

Although previous studies indicated redundant roles of BES/BZR family members in BR signaling (Yin et al., 2005; Lachowiec et al., 2018; Chen et al., 2019a; Chen et al., 2019b; Nolan et al., 2020), little has been reported about their functional differences. Here we found BEH3 as a non-canonical member of the BES/BZR family. *BEH3* is expressed preferentially in the vascular cambium, whereas *BES1* and *BZR1* expression is observed ubiquitously (Yin et al., 2002). Moreover, our results indicated the opposite function between BEH3 and BES1. In the context of *in vivo* vascular development, the gain-of-function *bes1-D* mutants accelerate cell differentiation of vascular stem cells and consequently decreased the number of (pro)cambial cell layer, eventually resulting in diminishment of the vascular size (Figure 2) (Kondo et al., 2014). Conversely, *BEH3* overexpression led to the increase in the number of (pro)cambial cell layer with enlargement of the vascular size. Consistently, the *beh3* loss-of-function mutants showed a decreased size of vascular area. Considering that BEH3 has a repressive effect on vascular cell differentiation in the VISUAL, these results suggest that BEH3 has a positive role in the maintenance of vascular stem cells probably by inhibiting excess cell differentiation (Figure 7D). Molecular analyses suggested that BEH3 retains a binding capability to the BRRE but loses a transcriptional repressor activity. Such a competitive action leads to the opposite behavior of BEH3. As a similar example in mammals, JunB, a transcription factor, exhibits weaker activity than its homolog, c-Jun, can repress c-Jun-mediated transactivation (Deng and Karin, 1993). In this study, our modeling and genetic analysis with the *beh3* mutants suggested that the presence of a weak or non-active type member in a transcription factor family enables a fine-tuned control of signaling outputs.

Taken together, we conclude that the competition among BES/BZR family members in the cambium potentially enables the stable and robust maintenance of vascular stem cells for ensuring continuous radial growth. However, it remains unclear why only BEH3 partially lacks the transcriptional repressor activity and why the competitive suppression by BEH3 is limited to the repressor activity. Further comparative interactome analysis between BEH3 and other BES1 homologs will uncover the mechanism underlying functional differences among BES/BZR proteins.

## METHODS

### Plant materials

Seeds of *Arabidopsis bes1-1* (SALK_098634), *beh1-1* (SAIL_40_D04), *beh3-1* (SALK_017577), and *beh4-1* (SAIL_750_F08) mutants were obtained from the Arabidopsis Biological Resource Center (ABRC), Ohio, USA. The *bzr1-1* single mutants and *bes1-1 bzr1-2* double mutants were generated previously using the CRISPR system (Saito et al., 2018), and *bes1-1 beh1-1, bes1-1 beh3-1, bes1-1 beh4-1, bes1-1 bzr1-2 beh3-1*, and *bes1-1 bzr1-2 beh4-1* mutants were generated by crossing. Chen et al. reported that the *bes1-1* mutants expresses one of the two splice variants of *BES1* (*BES1-S*); however, the other splice variant (*BES1-L*), which is functionally more important, is completely eliminated (Chen at al., 2019b, Jiang et al., 2015). The *bes1-1 bzr1-2 beh2-5 beh4-1* quadruple mutants were generated by the CRISPR/Cas9 system using pKAMA-ITACH vector (Tsutsui et al., 2017). The single guide RNA (sgRNA; Supplemental table 3.) sequence corresponding to the target gene, *BEH2*, was cloned into the *pKIR1*.*1* vector. The resulting construct was transformed into the *bes1-1 bzr1-2 beh4-1* triple mutants. In the T2 generation, *Cas9*-free plants with homozygous *beh2-5* mutation were selected as described previously (Tsutsui et al., 2017). The *bes1* quintuple mutants (*bes1-1 bzr1-2 beh1-2 beh2-5 beh4-1*) were also generated using the CRISPR/Cas9 system, which targeted the *BEH1* locus in *bes1-1 bzr1-2 beh2-5 beh4-1*. To generate the *bes1-1 beh3-2* double mutants and *bes1* hextuple mutants (*bes1-1 bzr1-2 beh1-2 beh2-5 beh3-3 beh4-1*), the *BEH3* locus was edited using the CRISPR/Cas9 system in *bes1-1* single mutants and *bes1* quintuple mutants, respectively. To generate a construct for *BEH3* overexpression under the control of a β-estradiol-inducible promoter, the coding sequence of *BEH3* (At4g18890.1) was amplified from Col-0 cDNA by PCR and then cloned into pENTR/D-TOPO (Life Technologies). Using LR clonase II (Life Technologies), the region between *attL1* and *attL2* in the entry vector was cloned into the destination vector, pER8-GW-CFP, which expressed the cyan fluorescent protein (CFP)-tagged proteins upon estrogen application (Ohashi-ito et al., 2010). To generate the *pBEH3:GUS* construct, a 1,997 bp fragment upstream of *BEH3* was amplified from Col-0 genomic DNA by PCR and cloned into pENTR/D-TOPO. The region between *attL1* and *attL2* in the entry vector was cloned into the destination vector, pGWB434. These constructs were transformed into Col-0 plants by the floral dip method (Clough et al., 1998). All genotypes used in this study were in the Columbia background. Plants were grown on conventional half-strength Murashige and Skoog (1/2 MS) agar plates (pH 5.7) at 22°C under continuous light, unless stated otherwise.

### Vascular cell Induction culture System Using Arabidopsis Leaves (VISUAL)

Plants were analysed using VISUAL as described previously (Kondo et al., 2016), with a slight modification. Briefly, cotyledons of 6-day-old seedlings grown in liquid 1/2 MS medium were cultured under continuous light in MS-based VISUAL induction medium containing 0.25 mg/l 2,4-D, 1.25 mg/l kinetin, and 20 µM bikinin. To observe ectopic xylem differentiation, a xylem indicator BF-170 (Sigma-Aldrich) was added to the induction medium (Nurani et al., 2020). At 4 days after induction, cotyledons were fixed in ethanol: acetic acid (3:1, v/v) and mounted with a clearing solution (chloral hydrate: glycerol: water = 8:1:2 [w/w/v]). Images obtained using a fluorescent stereomicroscope (M165, Laica) were binarized using the “Threshold” function of the ImageJ software (Schneider et al., 2012). Xylem differentiation ratio was calculated based on the BF-170 positive area cut-off by suitable thresholding per a whole cotyledon area.

### Microarray Experiments

To obtain the time-course maicroarray datasets, VISUAL was performed as described above. After VISUAL induction, cultured cotyledons (6 to 10 cotyledons) were collected every six hours. Total RNA was extracted from cultured cotyledons using an RNeasy Plant Mini Kit (Qiagen). Microarray experiments were performed using the Arabidopsis Gene 1.0STArray (Affymetrix) according to the standard Affymetrix protocol.

### Construction of the Coexpression Network

Based on the VISUAL time-course maicroarray data, the VISUAL-induced genes were defined as >8-fold changed genes compared with VISUAL for 0 and 72 h. Median normalized values in log_2_ scale of the VISUAL-induced genes (855 genes) were obtained from multiple transcriptome datasets: time-course, wild type versus *bes1-1* (Saito et al. 2018), wild type versus *apl* (Kondo et al. 2016), and SEOR1 cell-sorting data (Kondo et al. 2016). As described previously (Kondo et al., 2016), the VISUAL coexpression network was constructed using the weighted gene coexpression network analysis (WGCNA) package (Langfelder and Horvath, 2008). The adjacency matrix was calculated with soft thresholding power, which was chosen based on the criterion of scale-free topology (fit index =0.8).

### Confocal laser scanning microscopy

BF-170 signals in VISUAL samples were visualized using the FV1200 confocal laser scanning microscope (Olympus). Z-series fluorescence images were obtained at excitation and detection wavelengths of 473 and 490–540 nm, respectively. Z-projection images were created using the ImageJ software.

### Quantitative reverse transcription PCR (qRT-PCR)

Total RNA was extracted from cotyledons subjected to VISUAL experiments or whole seedlings in other experiments using RNeasy Plant Mini Kit (Qiagen). Quantitative PCR was performed using a LightCycler (Roche Diagnostics). *UBQ14* (for VISUAL experiments) and *ACT2* (for other experiments) were used as internal controls. Gene-specific primers and TaqMan Probe sets are listed in Supplemental Table 4.

### RNA-seq

Total RNA was extracted from cotyledons of Col-0 plants subjected to VISUAL experiments (Kondo et al., 2015) at 0, 30, and 72 h after induction using RNeasy Plant Mini Kit (Qiagen). RNA-seq analysis was carried out by Hokkaido System Science Co., Ltd.

### Expression and purification of BES/BZR family proteins

MBP-fused DNA-binding domains (DBDs) of BES1 (20T_103R) and BZR1 (21A_ 104R) were expressed and purified as described previously (Nosaki et al., 2018). The DBDs of BEH3 (4G_88S) and BEH4 (4G_89T) were cloned into pMAL-c2X (New England Biolabs) and subsequently transformed into *Escherichia coli* Rosetta (DE3) cells (Novagen). MBP-fused DBDs of BEH3 and BEH4 were expressed and purified in the same way as BES1 and BZR1.

### Analysis of BES/BZR proteins by size exclusion chromatography (SEC)

The purified proteins were loaded onto a Superdex 200 GL 10/300 (GE Healthcare) column and eluted with a buffer containing 20 mM Tris-HCl (pH 7.5), 250 mM NaCl, 1 mM DTT, and 5% glycerol. To estimate the multimerization state of BES/BZR family proteins, the following standards were used: thyroglobulin (*Mr* 670,000), bovine globulin (*Mr* 158,000), chicken ovalbumin (*Mr* 44,000), equine myoglobin (*Mr* 17,000), and vitamin B-12 (*Mr* 1,350) (Bio-Rad).

### EMSA

EMSAs of BES/BZR family proteins were performed as described previously (Nosaki et al., 2018).

### Luciferase (LUC) assay

Coding sequences (CDSs) of *BES1-L* (*At1g19350*.*3*), *BZR1* (*At1g75080*.*1*), and *BEH4* (*At1g78700*.*1*) were cloned and fused to the ligand-binding domain of the rat glucocorticoid receptor (GR). An approximately 1,952 bp sequence upstream of the *DWF4* transcription start site was cloned and fused to ELUC with the PEST domain (Tamaki et al., 2020) (Toyobo). *Rhizobium radiobacter* GV3101 MP90 strain harboring both effector and reporter constructs as well as a p19k suppressor construct was infiltrated into *Nicotiana benthamiana* leaves. After 2 days, leaf discs excised from the transformed leaves were incubated with 200 µM D-luciferin (Wako) for 2 h in white 24-well plates (PerkinElmer). Then, 10 µM dexamethasone (DEX) was added into each well, and LUC activity was measured using the TriStar2 LB942 (Berthold) system, as described previously (Tamaki et al., 2020).

### Histological analysis

Cross-sections of hypocotyls of seedlings were prepared as described previously (Kondo et al., 2014). Samples were fixed in FAA solution containing 40% (v/v) ethanol, 2.5% (v/v) acetic acid, and 2.5% (v/v) formalin. The fixed samples were gradually dehydrated using an ethanol dilution series and then embedded in Technovit 7100 resin (Kulzer). Sections of 5 μm thickness were prepared using the RM2165 microtome (Leica). Sliced samples were stained with 0.1% (w/v) toluidine blue for 1 min.

### Mathematical modeling

A mathematical model for the competitive binding between BEH3 and other homologs was constructed based on the assumptions that 1) there are *n* binding sites at the promoter, and 2) the weak-type BEH3 is competitively superior to the strong-type homologs in terms of binding to the promoter. The weak-type BEH3 occupies the vacant sites first, followed by the strong-type homologs, which occupy the remaining vacant sites. The binding probabilities of the weak-type BEH3 and strong-type homologs (*p*_*BEH*3_and *p*_0_, respectively) are proportional to the transcript abundance of *BEH3* and other homologs, respectively. Using *p*_*BEH*3_and *p*_0_, the probability that *i* BEH3 and *j* homologs bind to the promoter is given as the product of two binomial distributions as follows:

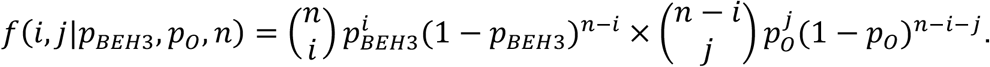

Binding of BEH3 and other homologs to the binding sites suppresses the transcription level of direct target genes leading to the repression of the proliferation of stem cells. The effect of the binding sites of BEH3 and other homologs on the cell proliferation rate of stem cells was designated as αand β, respectively. Because the effect of BEH3 is weaker than that of other homologs, α< β always holds. When *i* BEH3 and *j* homologs bind to the promoter, the cell proliferation rate is given as *α*^*i*^*β*^*j*^. The cell division rate should be proportional to the area inside the cambium, which was measured in experiments because the area inside the cambium increases with the increase in the rate of cell division. When there are multiple individuals, the expected rate of cell division of stem cell is calculated as follows:

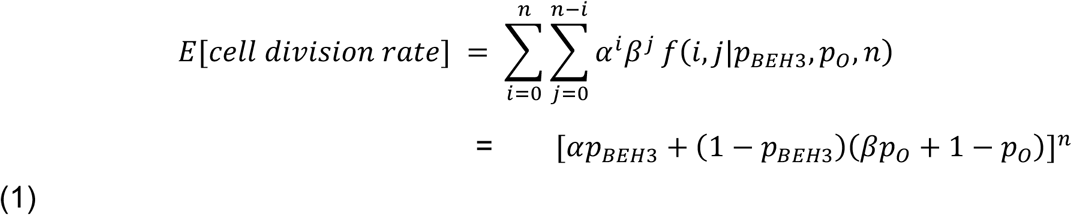

Using Equation (1), the variance of cell division rate can be calculated as:

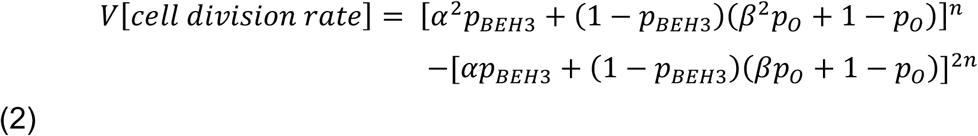

The coefficient of variation (CV) of the cell division rate was calculated as:

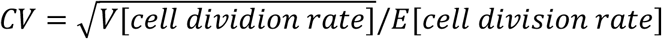

### Data availability

All data is available in the manuscript or the supplementary materials. All sequence reads were deposited in the NCBI Sequence Read Archive (SRA) database under the following accession numbers: XXXXXXX

## AUTHER CONTRIBUTIONS

T.F., M.S., and Y.K. designed the experiments, coordinated the project, and wrote the manuscript; T.F, M.S., H.U., S.S., W.Y., S.N., T.M., and Y.K. performed the experiments; A.S. performed mathematical model simulations; M.T. and H.F. participated in discussions. All authors reviewed and edited the manuscript.

## ACKNOWLEDGMENTS

We thank Yasuko Ozawa, Yuki Fukaya, and Akiho Suizu for technical support. We also thank Dr. Keiko U. Torii (University of Texas, Austin, Texas) for critically reading the manuscript and providing insightful comments. This work was funded by the Ministry of Education, Culture, Sports, Science and Technology, Japan (Scientific Research on Priority Areas and Scientific Research on Innovative Areas) (17H06476 and 20H05407 to Y.K., and 19H04855 to T.M.), and by the Japan Society for the Promotion of Science (17H05008 and 20K15815 to Y.K., 18H06065 and 20K15813 to T.F., 16H06377 to H.F., 19K23658 to S.N., and 18KK0170 and 18H02185 to W.Y.).

---

**Supplementary Figure 1.**
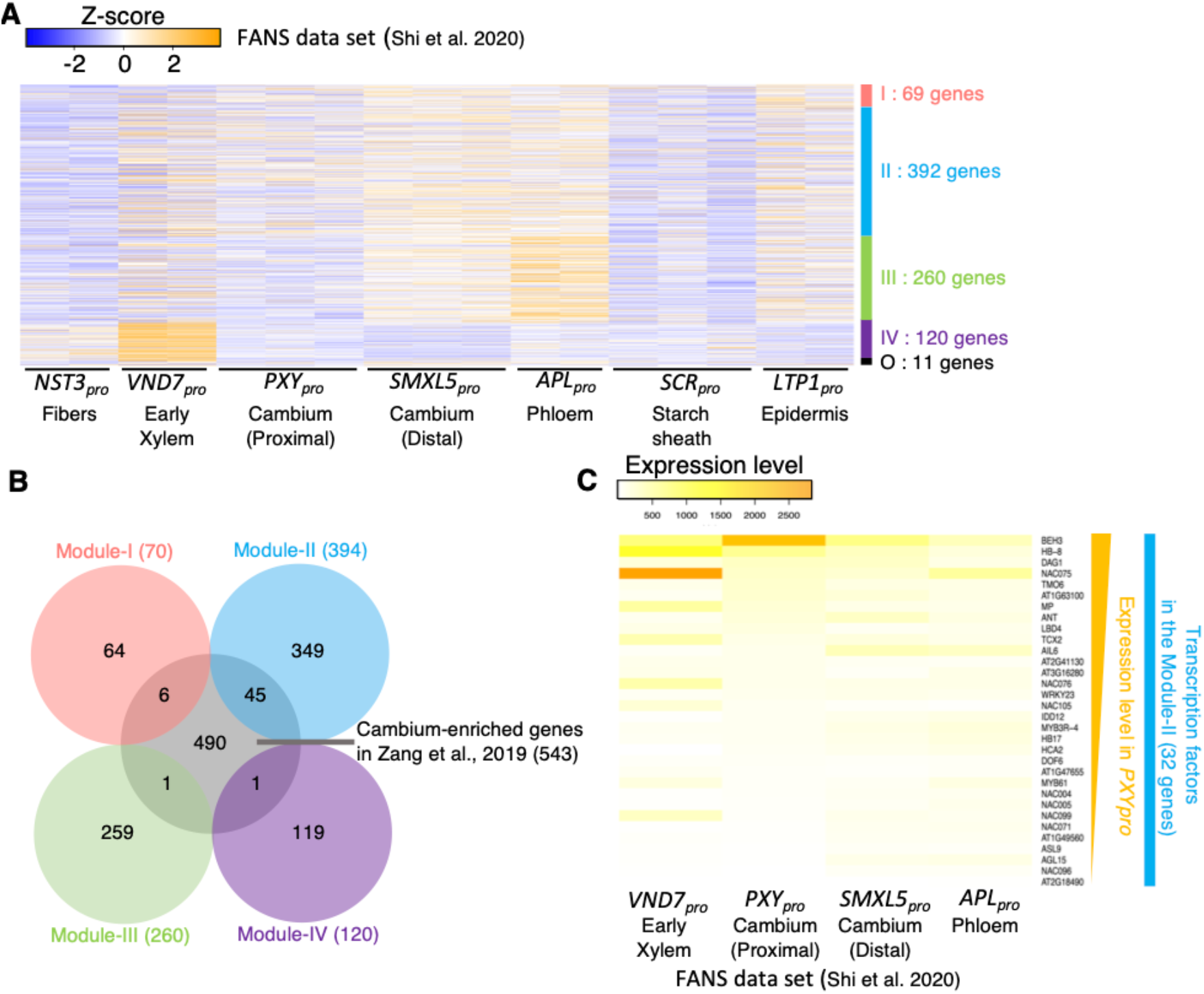
Comprehensive analysis of the VISUAL coexpression network and vascular-related transcriptome datasets. (Supports Figure 1) **(A)** Expression profiles of the VISUAL-induced genes in the FANS dataset (Shi et al., 2020). **(B)** Venn diagram of genes in each module of the VISUAL-induced genes and the cambium-enriched genes (Zang et al., 2019). **(C)** Expression levels of the cambium-related genes (Module-II) in the FANS dataset (Shi et al., 2020). The gene names are shown on the right side.

**Supplementary Figure 2.**
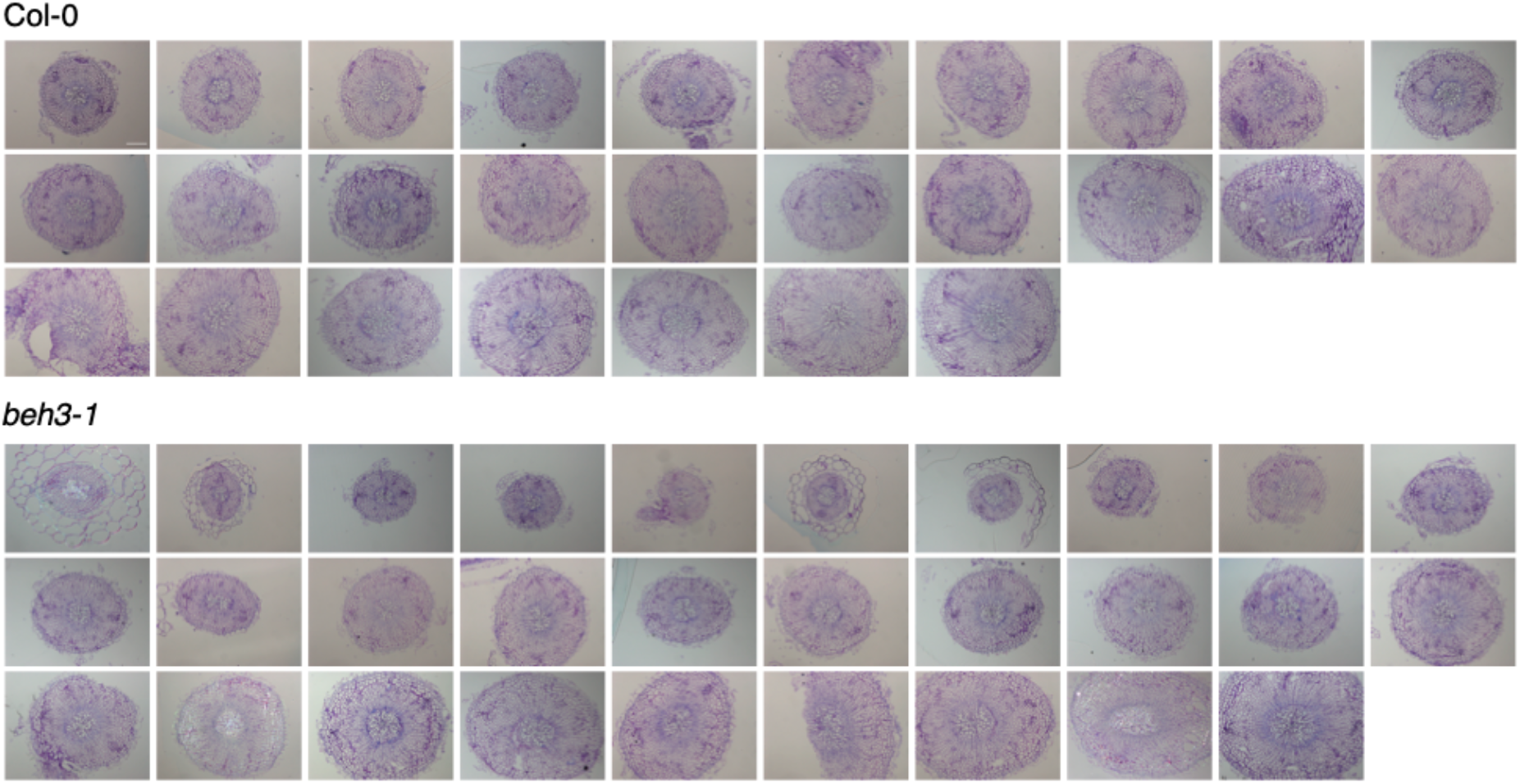
All section images of hypocotyl vasculature of WT and *beh3-1* mutants. (Supports Figure 2) All section images for Figure 2F. Sections of hypocotyl vasculature of WT and *beh3-1* mutants grown under short day conditions for 21 days. Scale bars = 100 µm.

**Supplementary Figure 3.**
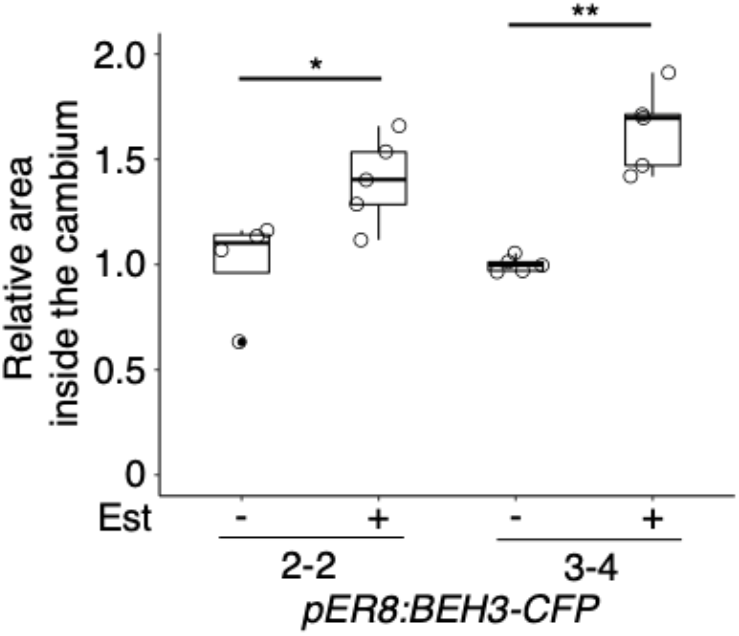
Sections of hypocotyl vasculature of the β-estradiol inducible *BEH3*-overexpressing plants. (Supports Figure 3) Plants were grown with or without 5 µM β-estradiol for 12 days. Relative areas inside the cambium were calculated in comparison with mean value of DMSO treated samples Quantified values were shown in box-and-whisker plots. White circles indicate the xylem differentiation ratio for each sample. (\**P* < 0.05 and \*\**P* < 0.01; Student’s *t*-test).

**Supplementary Figure 4.**
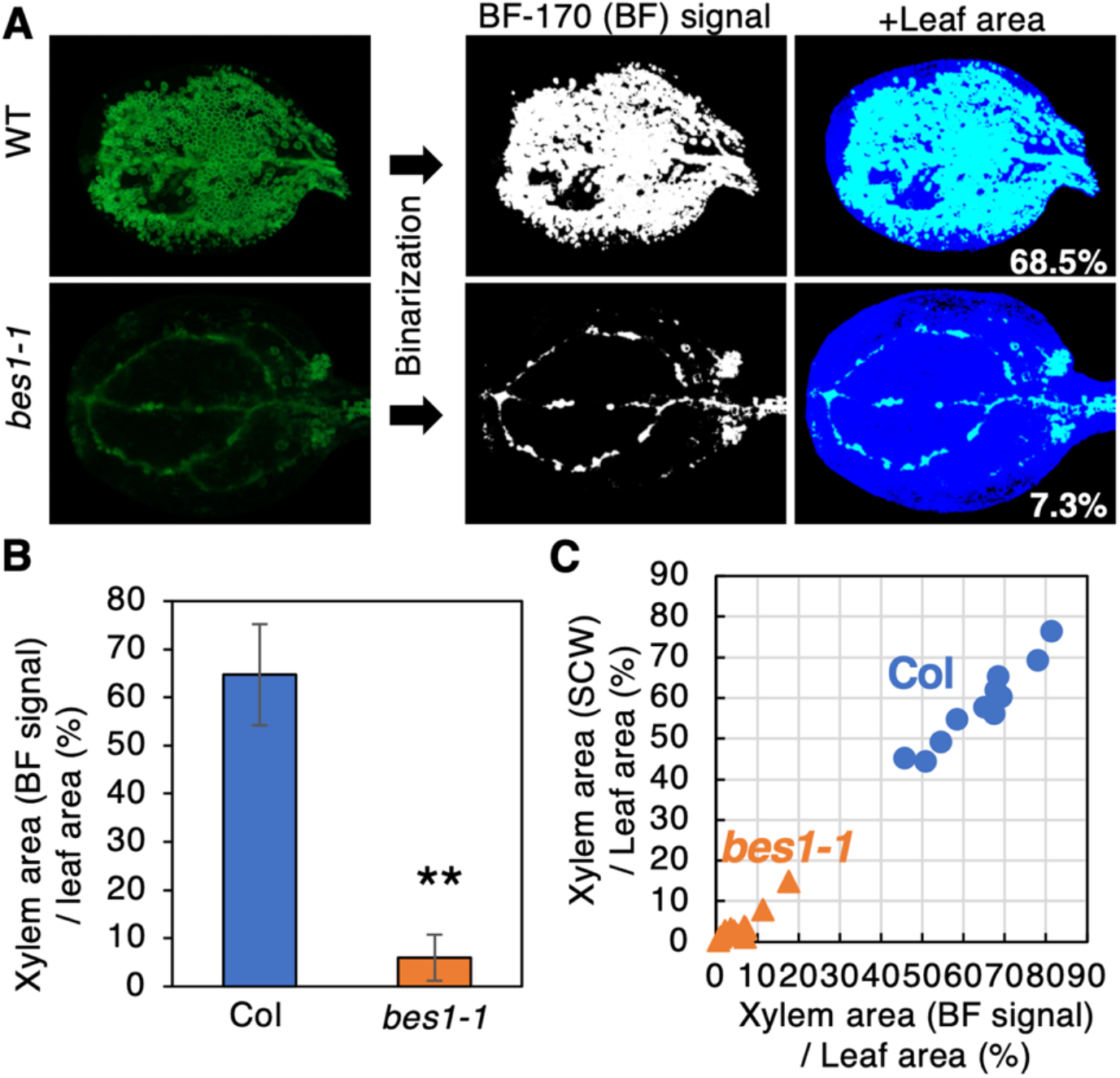
Semi-automatic calculation of xylem differentiation ratio by the binarization method. (Supports Figure 4) **(A)** The procedure of image thresholding in the WT and *bes1-1* after the VISUAL induction. The BF-170 signal (BF signal) area and the leaf area was shown by white and blue colour, respectively. The number written in the lower right represents the BF signal area per the leaf area (%) calculated by the binarization method. **(B)** Xylem area calculated based on BF signal per leaf area (%) in the WT and *bes1-1*. Statistical differences were examined by Student’s t-test (\*\**P* < 0.005). Error bars indicate SD (*n* = 12). **(C)** Comparison of two calculation methods for xylem differentiation ratio. X axis indicates the ratio of xylem area calculated from BF signal per the leaf area. Y axis indicates the ratio of xylem area calculated from bright field images per the leaf area. There is a strong positive correlation between them.

**Supplementary Figure 5.**
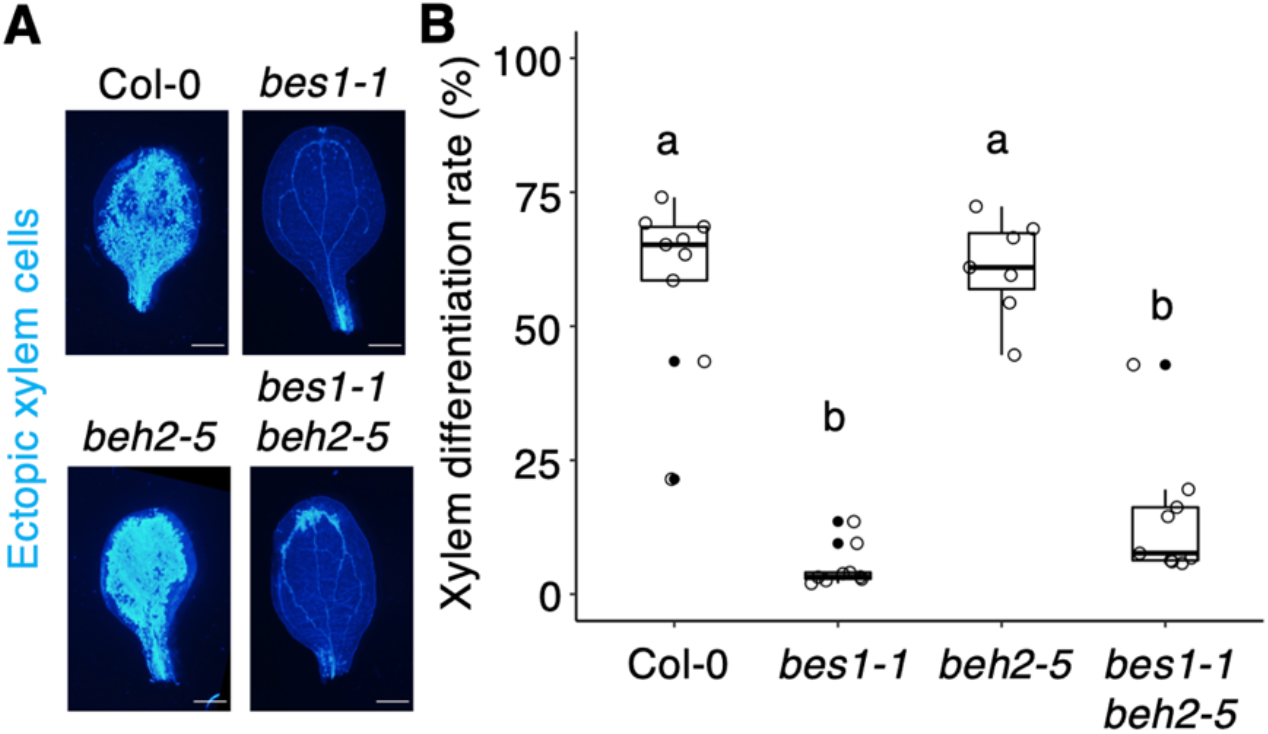
Ectopic xylem differentiation in VISUAL with *beh2* and *bes1 beh2* mutants. (Supports Figure 4) **(A)** A bright blue signal indicates xylem cell wall stained with BF-170. **(B)** Quantified xylem differentiation ratio was shown by box-and-whisker plots. White circles indicate the xylem differentiation ratio for each sample. Different letters indicate significant differences (*P* < 0.05; Tukey-Kramer test).

**Supplementary Figure 6.**
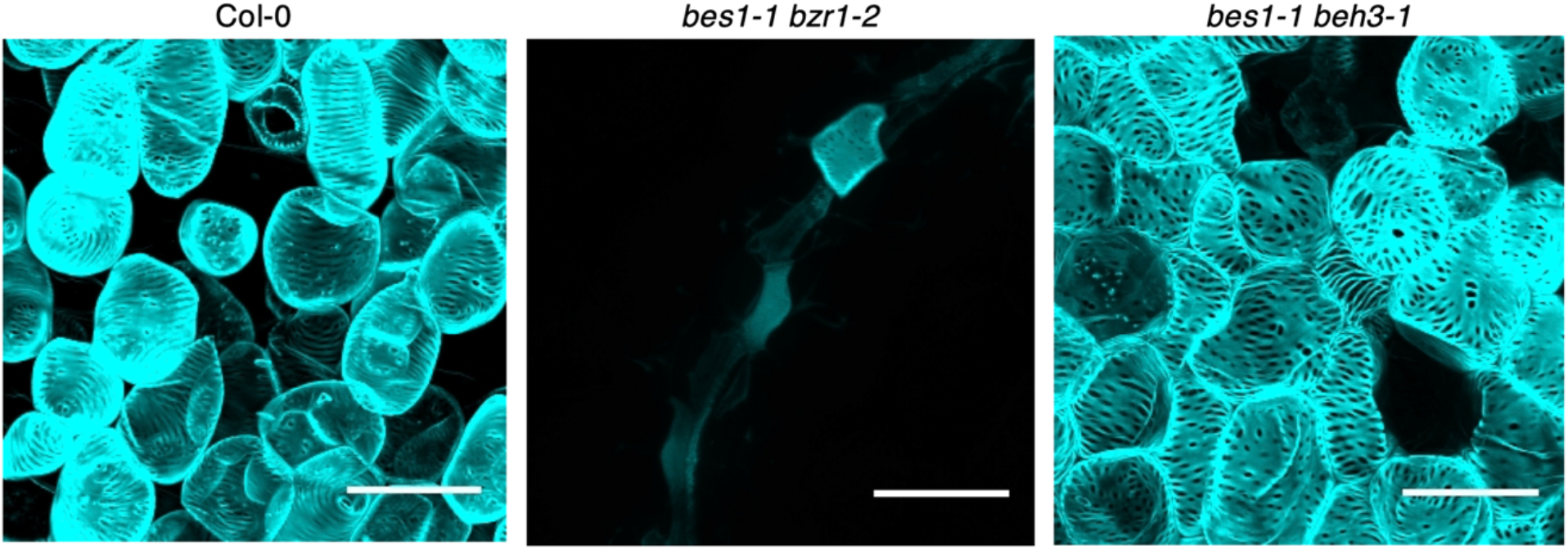
Ectopic xylem cells in VISUAL observed using confocal laser scanning microscope. (Supports Figure 4) Cotyledons of WT (Col-0), *bes1-1 bzr1-2*, and *bes1-1 beh3-1* were cultured with xylem indicator, BF170, for 4 days. A bright blue signal indicates BF170-incorporated xylem cell walls. Bars, 100 µm.

**Supplementary Figure 7.**
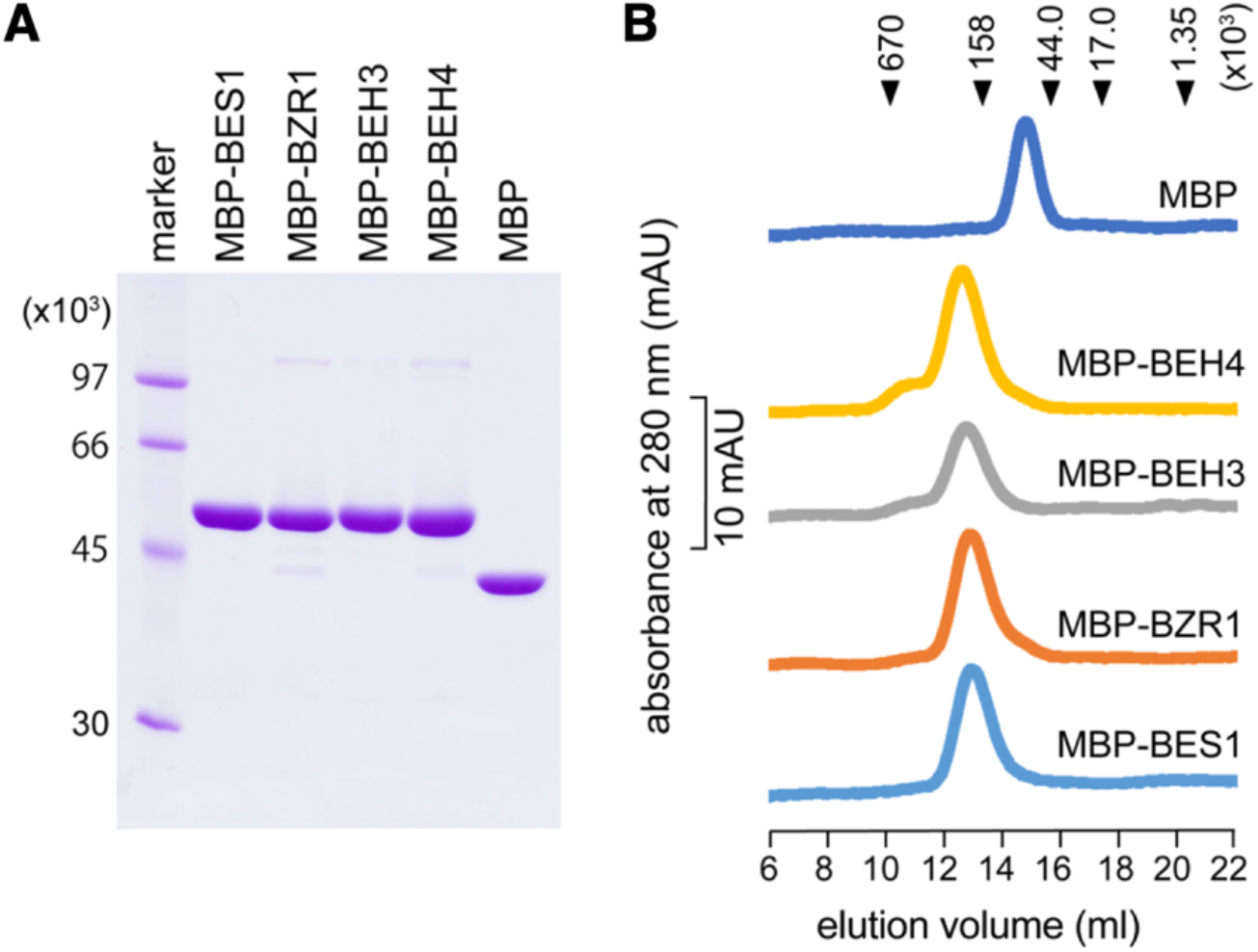
Preparation of MBP-fused proteins of BES/BZR family DBDs for EMSA assay. (Supports Figure 5) **(A)** Coomassie-stained SDS-PAGE analysis of the maltose-binding protein (MBP)-fused BES1, BZR1, BEH3 and BEH4 DBDs after purification. **(B)** Oligomeric state analysis of MBP-fused BES1, BZR1, BEH3 and BEH4 DBDs by size exclusion chromatography (SEC). The peak positions of the marker proteins are indicated by the black triangles at the top of the chromatogram.

**Supplementary Figure 8.**
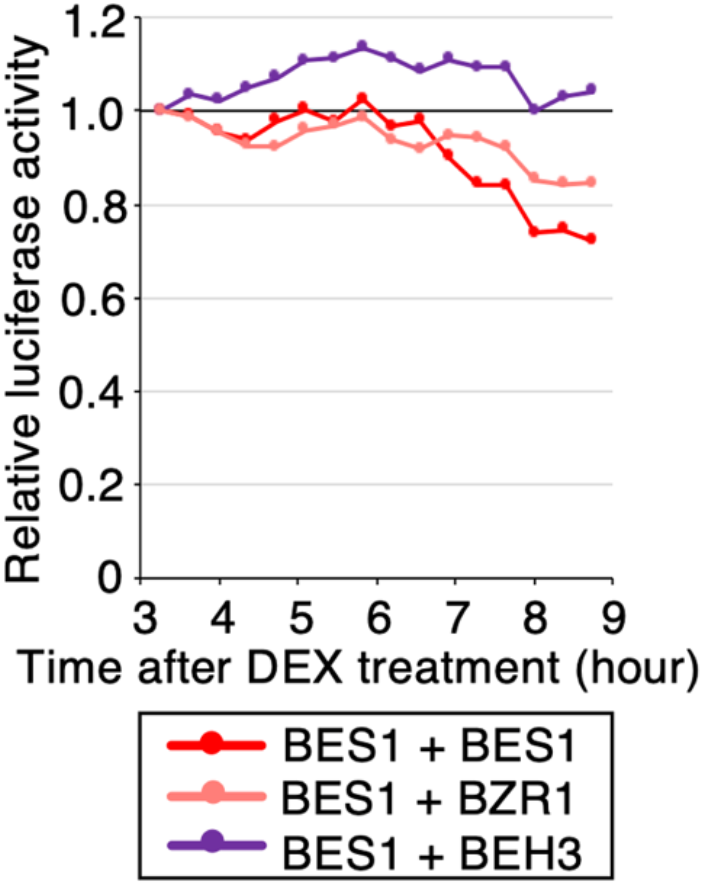
Combinatory effects on transcriptional repressor activity in the *Nicotiana benthamiana* transient expression assay. (Supports Figure 5) When *BZR1* or *BEH3* was co-expressed with *BES1*, luciferase (LUC) activities in *pDWF4:ELUC* transgenic plants were calculated as average photon counts per second of six leaf disks and were normalized relative to values of DEX-treated samples harbouring an empty effector construct.

**Supplementary Figure 9.**
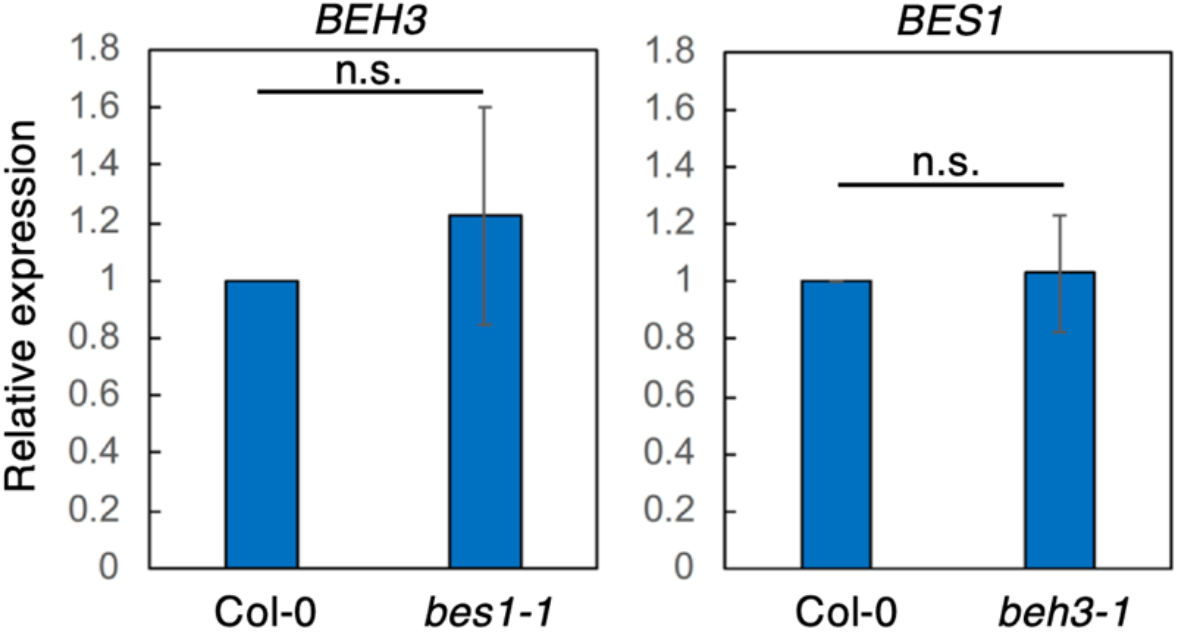
Expression levels of *BEH3* in *bes1-1* and *BES1* in *beh3-1* at 72 hours after VISUAL induction. (Supports Figure 4) The data are indicated as relative fold change to Col-0 (\**P*<0.05, \*\**P*<0.01; Student’s t-test, *n* = 3, error bar = SD).

**Supplementary Figure 10.**
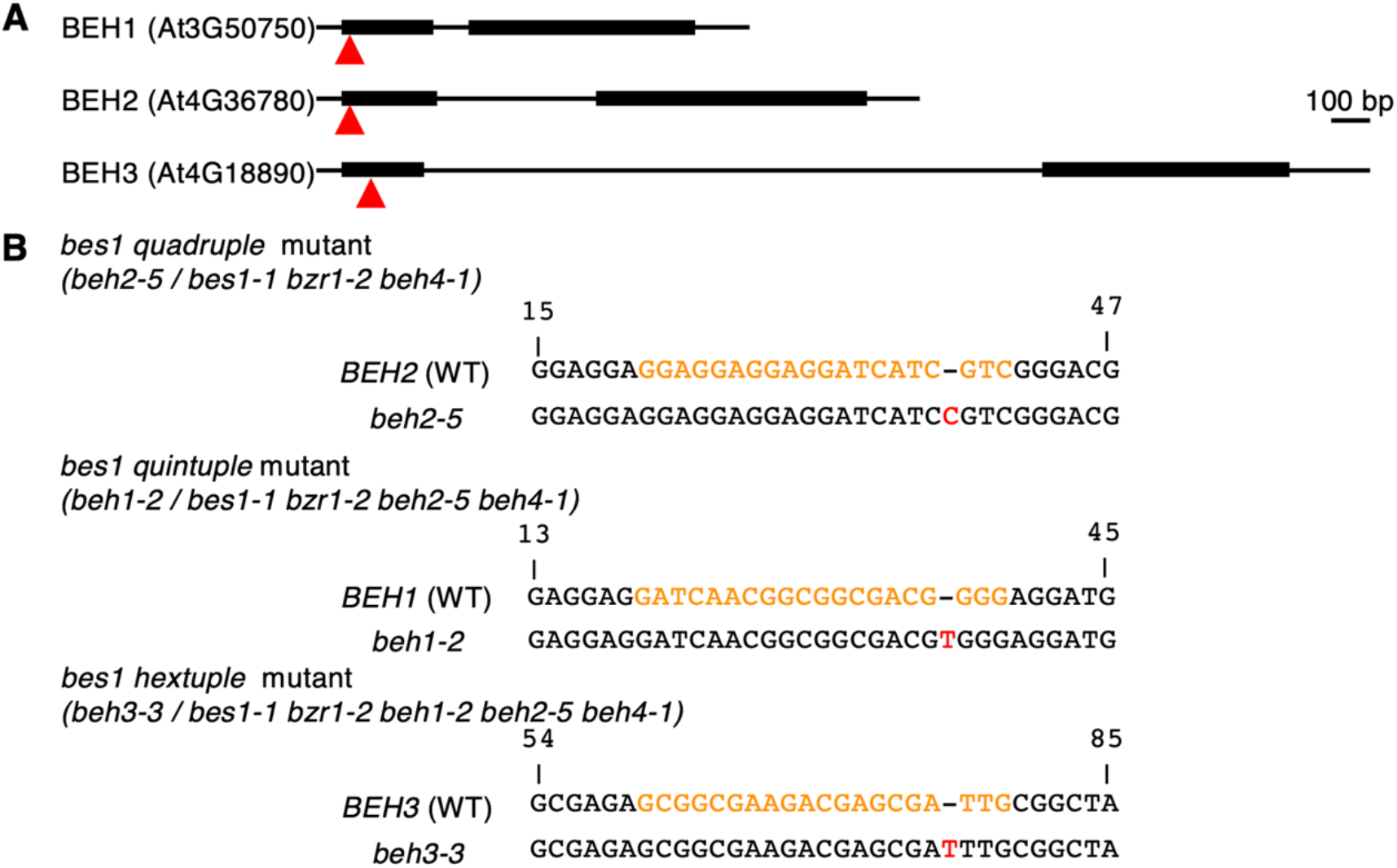
Construction of genome-edited lines for BES/BZR homologs. (Supports Figure 6) **(A)** Genomic structure of the encoding region of BEH1, BEH2, and BEH3. Red arrowheads indicate location of target site. **(B)** The targeted sequence (orange) and a part of sequence in both wild-type line and mutant lines are shown. Each genome-edited line has a 1-bp insertion (red).

**Supplementary Figure 11.**
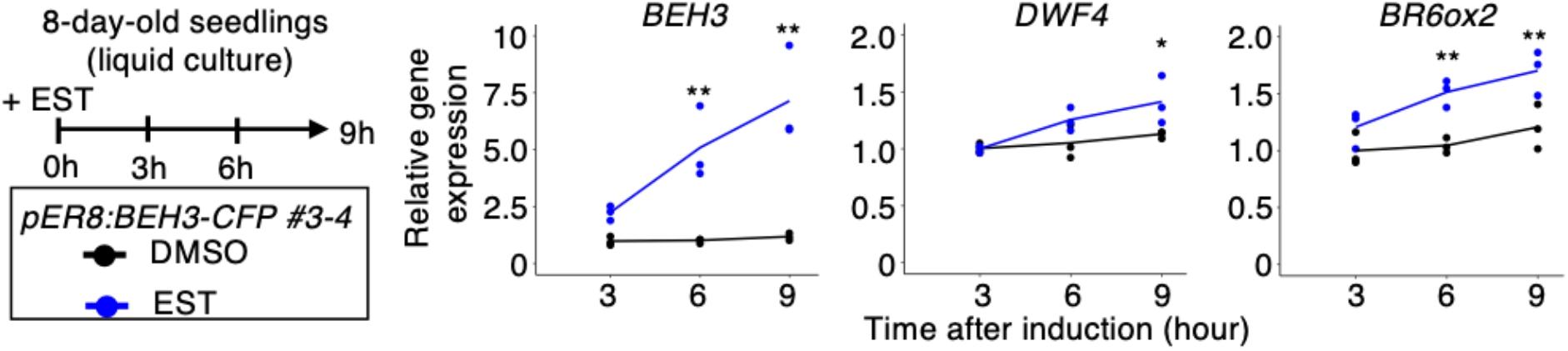
Time-course expression analysis of BES/BZR target genes in the β-estradiol-inducible *BEH3* overexpression line. (Supports Figure 6) Eight-day-old seedlings were treated with 10 µM β-estradiol to induce *BEH3* expression. Relative gene expression levels were calculated in comparison with the DMSO-treated sample at 3 h after induction (\**P* < 0.05, \*\**P* < 0.01; two-way ANOVA; *n* = 3). Values of statistical analysis listed in Supplemental Table 2.

**Supplementary table 1.**
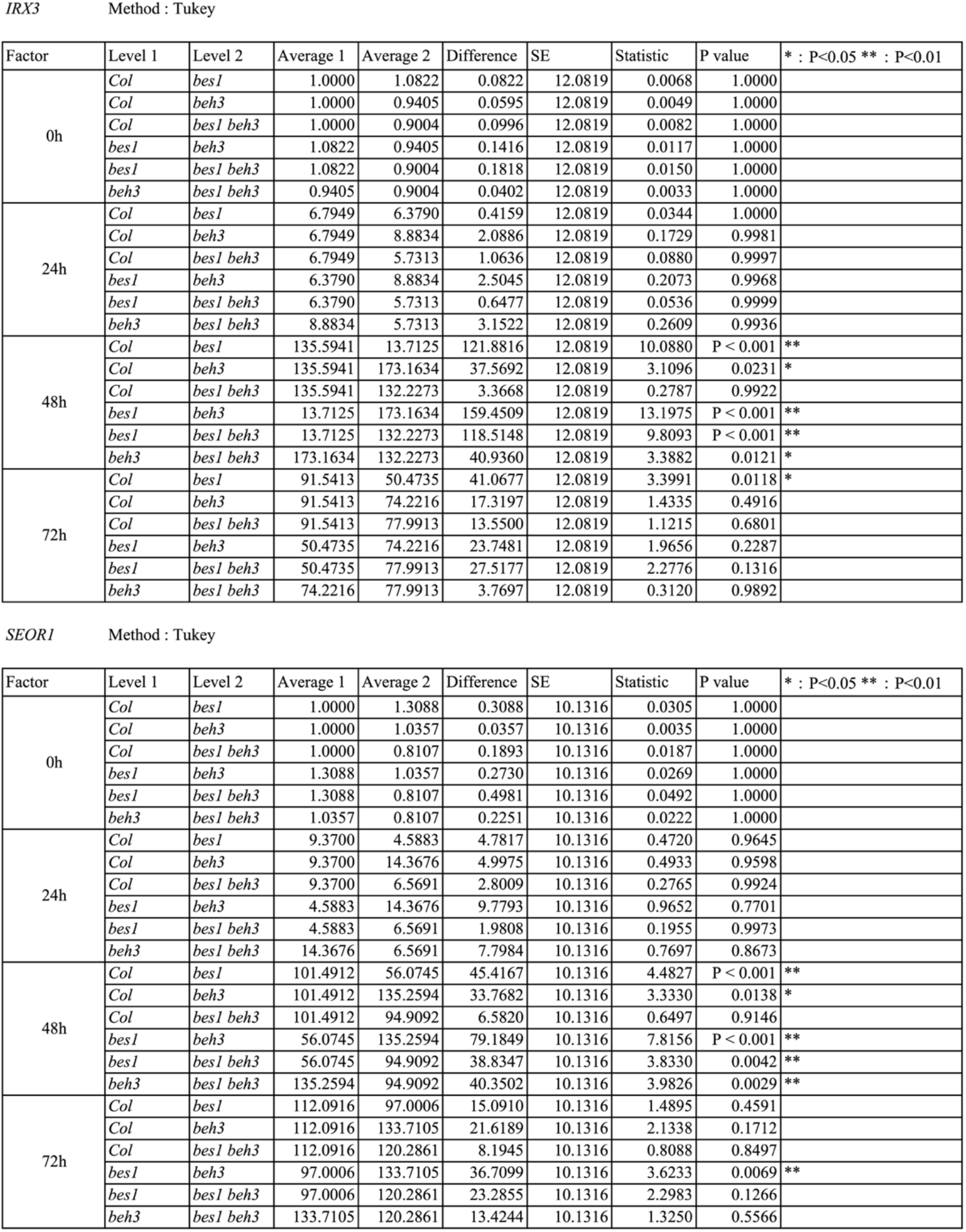
Results of two-way analysis of variance (ANOVA) for Figure 4E.

**Supplementary table 2.** Results of two-way analysis of variance (ANOVA) for Supplemental Figure 11.

*BEH3* Method : Tukey
| Factor | Level 1 | Level 2 | Average 1 | Average 2 | Difference | SE | Statistic | P value | * : P<0.05 ** : P<0.01 |
| --- | --- | --- | --- | --- | --- | --- | --- | --- | --- |
| 3h | DMSO | Est | 1.0000 | 2.2876 | 1.2876 | 0.9152 | 1.4068 | 0.1848 |  |
| 6h | DMSO | Est | 1.0340 | 5.1796 | 4.1456 | 0.9152 | 4.5295 | P < 0.001 | ** |
| 9h | DMSO | Est | 1.2027 | 7.2724 | 6.0697 | 0.9152 | 6.6318 | P < 0.001 | ** |

*BR6ox2* Method : Tukey
| Factor | Level 1 | Level 2 | Average 1 | Average 2 | Difference | SE | Statistic | P value | * : P<0.05 ** : P<0.01 |
| --- | --- | --- | --- | --- | --- | --- | --- | --- | --- |
| 3h | DMSO | Est | 1.0000 | 1.2068 | 0.2068 | 0.1267 | 1.6316 | 0.1287 |  |
| 6h | DMSO | Est | 1.0450 | 1.5146 | 0.4696 | 0.1267 | 3.7054 | 0.0030 | ** |
| 9h | DMSO | Est | 1.2062 | 1.7024 | 0.4961 | 0.1267 | 3.9144 | 0.0021 | ** |

*DWF4* Method : Tukey
| Factor | Level 1 | Level 2 | Average 1 | Average 2 | Difference | SE | Statistic | P value | * : P<0.05 ** : P<0.01 |
| --- | --- | --- | --- | --- | --- | --- | --- | --- | --- |
| 3h | DMSO | Est | 1.0000 | 0.9968 | 0.0032 | 0.0937 | 0.0340 | 0.9734 |  |
| 6h | DMSO | Est | 1.0486 | 1.2521 | 0.2035 | 0.0937 | 2.1705 | 0.0507 |  |
| 9h | DMSO | Est | 1.1264 | 1.4091 | 0.2827 | 0.0937 | 3.0160 | 0.0107 | * |

**Supplementary table 3.**
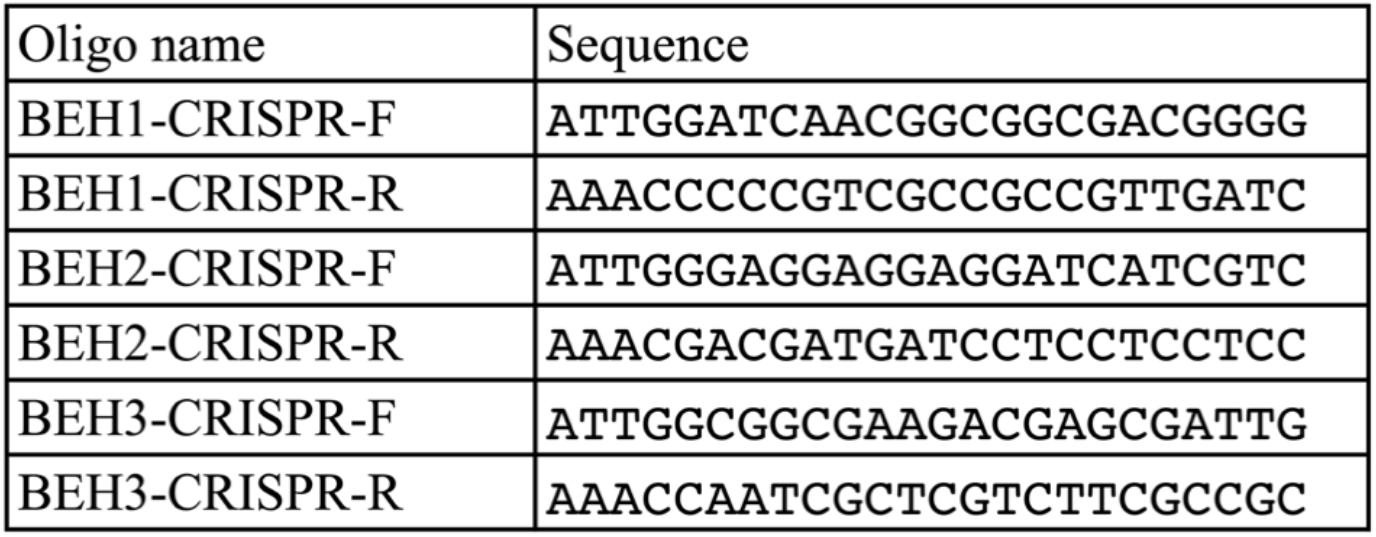
Sequences of sgRNA for CRISPR/Cas9 system.

**Supplementary table 4.**
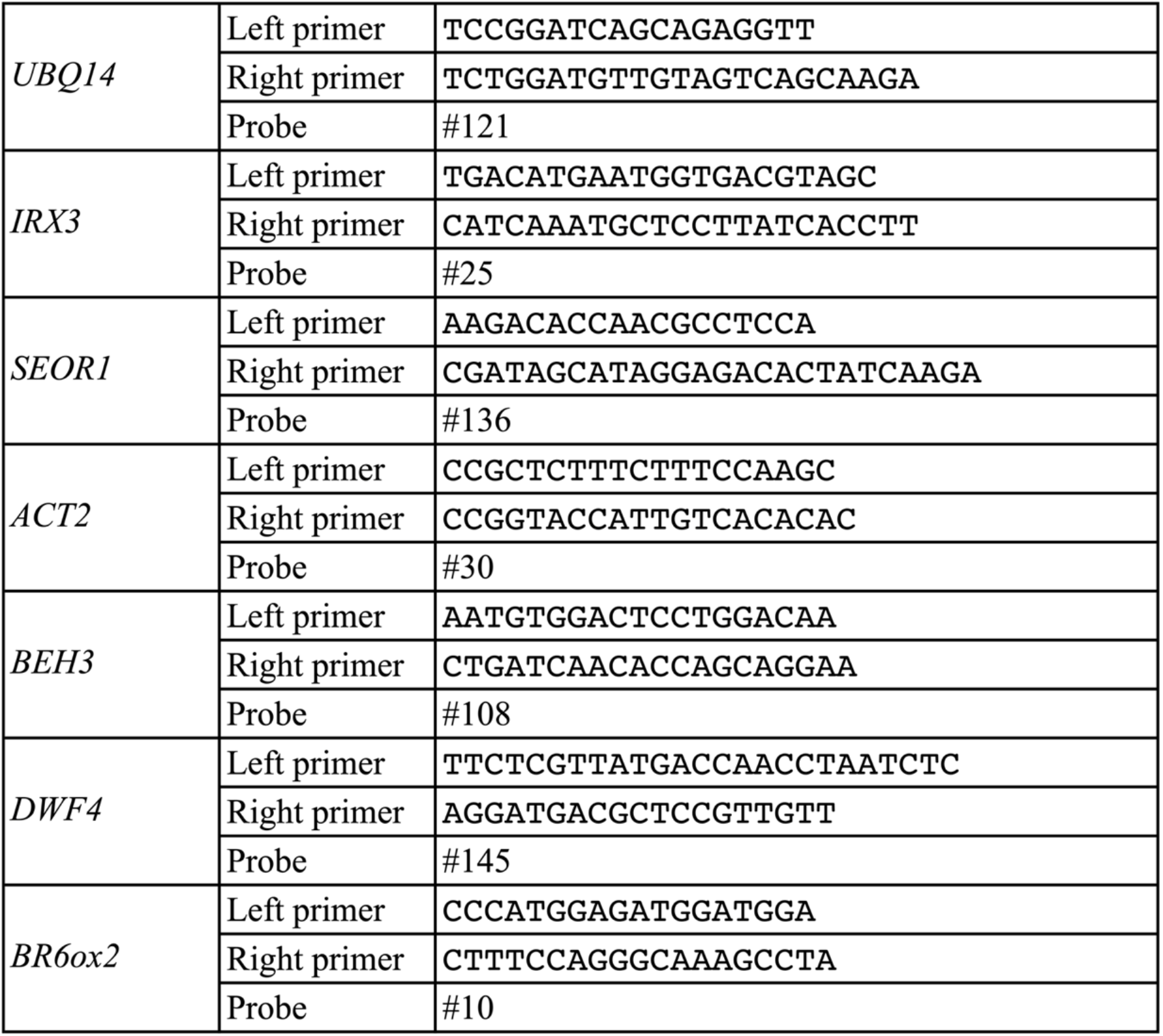
List of qRT-PCR primers.

